# Protein Moonlighting Revealed by Non-Catalytic Phenotypes of Yeast Enzymes

**DOI:** 10.1101/211755

**Authors:** Adriana Espinosa-Cantú, Diana Ascencio, Selene Herrera-Basurto, Jiewei Xu, Assen Roguev, Nevan J. Krogan, Alexander DeLuna

## Abstract

A single gene can partake in several biological processes, and therefore gene deletions can lead to different—sometimes unexpected—phenotypes. However, it is not always clear whether such pleiotropy reflects the loss of a unique molecular activity involved in different processes or the loss of a multifunctional protein. Here, using *Saccharomyces cerevisiae* metabolism as a model, we systematically test the null hypothesis that enzyme phenotypes depend on a single annotated molecular function, namely their catalysis. We screened a set of carefully selected genes by quantifying the contribution of catalysis to gene-deletion phenotypes under different environmental conditions. While most phenotypes were explained by loss of catalysis, 30% could be readily complemented by a catalytically-inactive enzyme. Such non-catalytic phenotypes were frequent in the Alt1 and Bat2 transaminases and in the isoleucine/valine-biosynthetic enzymes Ilv1 and Ilv2, suggesting novel "moonlighting" activities in these proteins. Furthermore, differential genetic-interaction profiles of gene-deletion and catalytic mutants indicated that *ILV1* is functionally associated to regulatory processes, specifically to chromatin modification. Our systematic study shows that gene-loss phenotypes and their genetic interactions are frequently not driven by the loss of an annotated catalytic function, underscoring the moonlighting nature of cellular metabolism.

## INTRODUCTION

The phenotypic impact of a mutation is amongst the most useful genetics tool providing insights into genes' functions in biological systems. Functional genomics has produced vast amounts of phenotypic data in different model organisms from yeast to mammals (Pollard 2003; Shalem *et al.* 2015; Norman and Kumar 2016). Still, deciphering the relationship between genotype and phenotype is a central challenge in system genetics (Baliga *et al.* 2017). Screens based on gene-deletion or RNAi perturbations have shown that many genes are associated to multiple and sometimes unexpected traits (Dudley *et al.* 2005; Ohya *et al.* 2005; Hillenmeyer *et al.* 2008; Zou *et al.* 2008; Carter 2013; Deutschbauer *et al.* 2014). A deep understanding of the molecular bases of such genetic pleiotropy contributes to our understanding of gene evolution, development, and disease (Promislow 2004; Wagner and Zhang 2011; Guillaume and Otto 2012; Hill and Zhang 2012; Smith 2016; Ittisoponpisan *et al.* 2017).

Genetic pleiotropy resulting from protein depletion may arise either from the loss of a single molecular function that impacts many cellular processes, or from the loss of more than one molecular function carried out by a single polypeptide (Hodgkin 1998; Stearns 2010; Paaby and Rockman 2013). Proteins were originally considered to be monofunctional, with a single highly-specific molecular function as proposed in the key and lock model (discussed in (Piatigorsky 2007; Copley 2012)). Nowadays, different mechanisms of protein multifunctionality have been described (Kirschner and Bisswanger 1976; Copley 2003; Khersonsky *et al.* 2006).

Moonlighting proteins are defined as polypeptides with two or more independent molecular activities, which are not the result of gene fusion events (Jeffery 1999; Piatigorsky 2007). Such proteins can provide selective advantage to an organism and are hotbeds for the evolution of molecular function (Jensen 1976; Piatigorsky 2007; Espinosa-Cantú *et al.* 2015; Jeffery 2015). In moonlighting enzymes, proteins exert other molecular functions (*e.g*. structural scaffold, transcription factor) independently of their catalytic activity. Indeed, the moonlighting behavior of enzymes is usually confirmed in mutants that lack catalysis but retain any other molecular activity. Protein moonlighting gives rise to molecules that link metabolism with regulation of gene expression (Shi and Shi 2004; Commichau and Stülke 2008; Boukouris *et al.* 2016), cellular crosstalk (Entelis *et al.* 2006; Hill *et al.* 2013; Torres-Machorro *et al.* 2015), pathogenesis, and disease (Yoshida *et al.* 2001; Sriram *et al.* 2005; Henderson and Martin 2013; Zanzoni *et al.* 2015).

Over 300 examples of moonlighting proteins have been characterized in Archaea, Prokaryotes, and Eukaryotes, from model to non-model organisms (Hernández *et al.* 2014; Mani *et al.* 2015). In the budding yeast *Saccharomyces cerevisiae*, over 30 moonlighting enzymes have been characterized, many of which are enzymes of core metabolic pathways (reviewed in (Gancedo and Flores 2008)). Some moonlighting proteins have been identified by function-specific screens (Zelenaya-Troitskaya *et al.* 1995; Hall *et al.* 2004; Chen *et al.* 2005; Scherrer *et al.* 2010). More recent studies have aimed to identify moonlighting proteins by computational approaches (Chapple *et al.* 2015; Hernández *et al.* 2015; Pritykin *et al.* 2015; Khan and Kihara 2016). Still, systematic strategies to directly identify protein moonlighting are currently missing, and most known examples have been recognized by chance. Hence, the question remains of how frequent the moonlighting phenomenon is, and what are the underlying molecular mechanisms (Gancedo and Flores 2008; Huberts and van der Klei 2010; Khersonsky and Tawfik 2010; Hernández *et al.* 2014).

Here, we asked if annotated molecular activities are enough to explain the pleiotropic behavior of genes in yeast. We focused on enzymes from the amino acid biosynthesis metabolism and quantified the contribution of catalysis to cellular phenotypes. To do so, we systematically compared the growth phenotypes of gene-deletion (knockout) and site-directed loss-of-catalysis (catalytic) mutants in different growth conditions; high-quality data was obtained for eleven enzymes in our screen. While catalytic mutants recapitulated most gene-deletion phenotypes, we found consistent phenotypes of several enzymes that did not depend on their annotated catalytic activity, suggesting moonlighting functions. We further explored the genetic-interaction landscape of *ILV1* and showed that catalysis-independent genetic interactions fell into discrete functional groups. In doing so, we shed light on the cellular processes in which Ilv1 is involved independently of its threonine deaminase activity.

## MATERIALS AND METHODS

### Strains, plasmids, and media

The parental *S. cerevisiae* strain used for gene replacement was Y8205 (MAT-α *can1*∆::STE2pr-Sp_his5 *lyp1*∆::STE3pr-LEU2 *his3*∆ *leu2*∆ *ura3*∆). Because strains from the yeast deletion collection may bear non-linked mutations and aneuploidies (Hughes *et al.* 2000), all gene knockouts were generated de novo on an intact isogenic parental background. For gene replacement, the *nat-* or *hph*-resistance cassettes were amplified from pAG25 or pAG32, respectively. Phenotypic complementation of enzyme-coding genes included in the phenotypic screen (*GENE*_*i*_) was done by transforming knockout strains (Y8205 *gene*_*i*_∆::*nat*) with centromeric plasmids bearing (1) the intact *GENE*_*i*_ sequence (resulting in strain WT_*i*_), (2) the *gene*_*i*_ catalytic mutant (CM_*i*_ strain), or (3) the empty plasmid (KO_*i*_ strain). Centromeric low-copy number plasmids were from the MoBY-ORF collection of open reading frames from *S. cerevisiae* with their corresponding 5’-promoter and 3’-UTR sequences (Ho *et al.* 2009). The empty plasmid was p5586 MoBY with no yeast-gene sequence. Strain Y8205 *his3*∆::*nat* + p5586 was used as the "WT" reference (WT_*ref*_). For all *GENE*_*i*_ that have a *GENE_i_'* paralog, plasmid transformations were also done on the Y8205 *gene*_*i*_∆::*nat gene_i_'*∆::*hph* double-knockout background. Primer sequences for gene-knockout and confirmation PCR were obtained from the Yeast Deletion Project (http://sequence-www.stanford.edu/group/yeast_deletion_project/deletions3.html). All strains used in this study are shown in **Table S1**.

Rich medium was Yeast-extract Peptone Dextrose (YPD); Synthetic-complete medium (SC) was 6.7 g/L yeast nitrogen base without amino acids and 2% glucose (unless otherwise indicated) supplemented with 0.2% drop-out amino acid supplement mix (Cold Spring Harbor Laboratory Manual 2005); see **Table 1** for drop-out media used for auxotrophy tests. Antibiotic-resistant strains were grown in media supplemented with 100 μg/ml nourseothricin (ClonNAT, Werner BioAgents), 200 μg/ml G418 (Invitrogen), or 200 μg/ml hygromycin (Invitrogen); ammonium was replaced by 1g/L monosodium glutamic acid in SC medium with antibiotics.

**Table 1.**
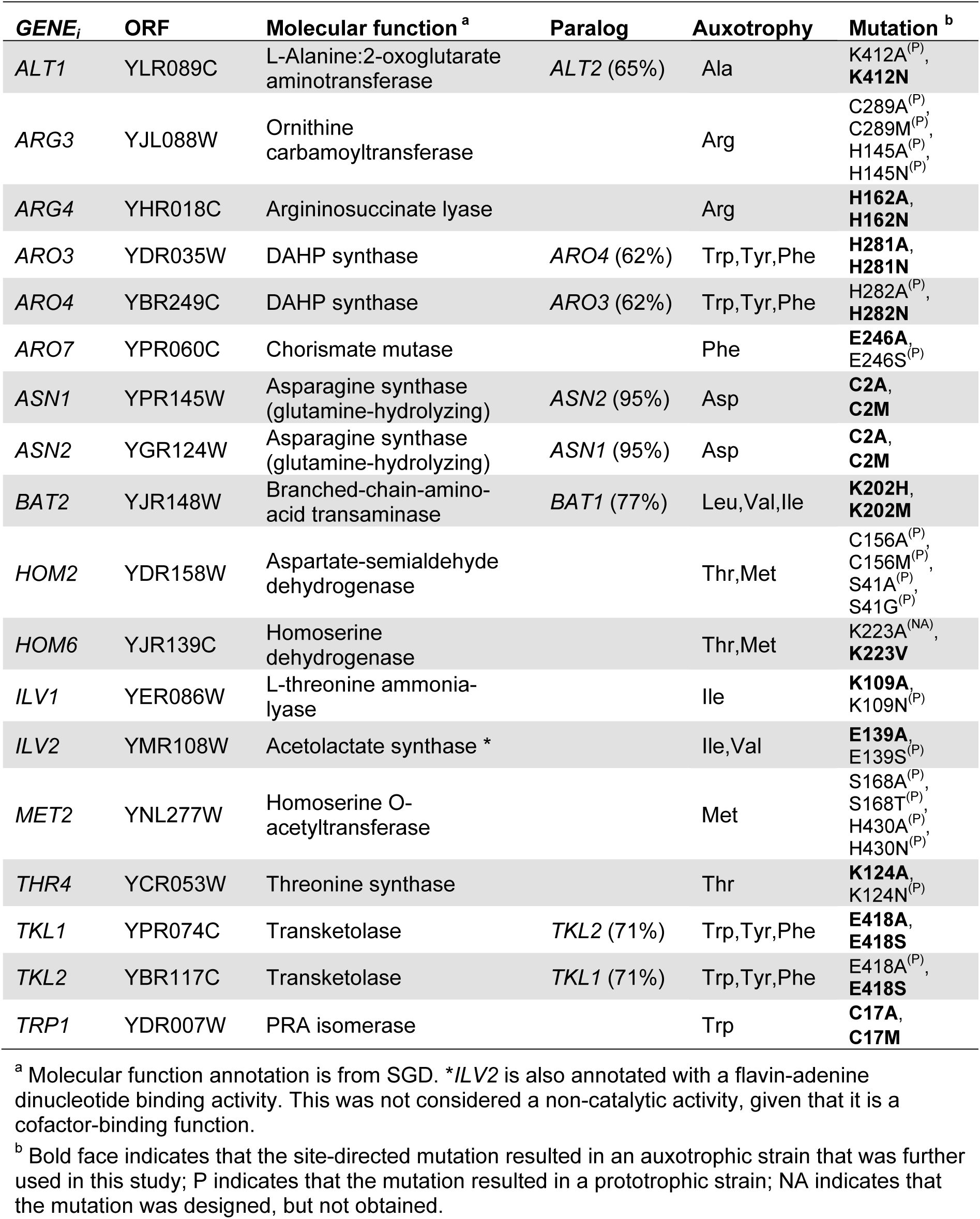
Amino acid biosynthesis enzymes and catalytic mutants in this study.

### Catalytic-mutant design and site-directed mutagenesis

To generate yeast strains expressing enzymes with no catalytic activity, a single catalytic residue for each *GENE*_*i*_ documented in the Catalytic Site Atlas (Furnham *et al.* 2014) or the MACiE Database (Holliday *et al.* 2012) was targeted (**Table S2**). Mutations were directed to an amino acid involved in the early stages of the reaction to prevent incomplete catalytic inactivation that could result in neomorphic phenotypes. If required, a second round of mutagenesis was directed to another residue involved in catalysis or in cofactor binding. Once a catalytic residue was selected, its conservation between the repository PDB and the query gene was established by sequence and structural alignments in PyMOL. In queries without PDB, models were obtained from the Swiss Model repository or generated in automatic (Bienert *et al.* 2017). For each *GENE*_*i*_, we substituted the catalytic residues to a residue with similar physicochemical characteristics, or to alanine.

Site-directed replacements of plasmid inserts were introduced by PCR with ∼40 base-pair of overlapping primers with the mutation (**Table S3**). PCR was conducted in two stages: the first stage included two independent reactions with the mutagenic forward or reverse primer in a final volume of 25 μl each (conditions below). PCR reactions were done with 5 ng/μl plasmid in a final mix of 25 μl with 1U of Phusion polymerase (Fermentas), 1x high fidelity buffer, 0.2 μM dNTPs, and 0.5 μM primers. PCRs were first incubated 3 minutes at 98°C; then 15 cycles of 98°C for 30 seconds, 55°C for 25 seconds and 7.5 minutes at 72°C, with a final extension of 10 minutes at 72°C. The second stage were 50 μl mixes of the two PCRs from the first stage, using the same parameters, and followed by an overnight incubation at 37°C with 0.5 μl of Dpn1 20,000 U/ml (Biolabs) to deplete for original non-mutagenized plasmid sequences. A 5 μl sample of the digested PCR product was used to transform calcium-competent *Escherichia coli* BUN20 (Li and Elledge 2005). Selection was done in LB 5 μg/ml tetracycline, 100 μg/ml kanamycin, and 12.5 μg/ml chloramphenicol. Plasmids from at least three clones from each transformation were sequenced with confirmation primers (**Table S3**). Confirmed mutated plasmids were used to transform yeast by lithium acetate (Schiestl and Gietz 1989), with selection for uracil prototrophy or geneticin resistance.

For amino acid auxotrophy confirmation by drop-spot assays, strains were grown for 36 hrs in SC-uracil, cultures were diluted to OD_600_=0.5, and 3 μl of 10^0^-10^6^ dilutions were inoculated onto SC-aa and SC-uracil plates. Cultures were incubated at 30°C during 72‒96 hrs and images were taken at two different time points for each set of strains.

### Automated phenotypic characterization

Plasmid-transformed strains were inoculated into 100 μl of YPD with antibiotics in 96-well microtiter plates (Corning 3585) and grown without shaking at 30°C. Saturated cultures (5 μl) were inoculated into 150 μl of different growth media for phenotypic analysis under different conditions (see below). Growth was monitored in an automated robotic system (Tecan Freedom EVO200) that integrates a plate carrousel (Liconic STX110), a plate reader (Tecan Infinite M1000), an orbital plate shaker, and a robotic manipulator arm. The equipment was maintained in an environmental room at constant temperature (30°C) and relative humidity (70%). Absorbance at 600 nm (OD_600_) was measured every 60-90 min after vigorous shaking and growth kinetics were followed for 24-30 hours. Unless otherwise noted, all experiments were performed in five independent biological replicates result of individual colonies from plasmid transformation.

Phenotypic characterizations were done under different growth conditions (C_*j*_): YPD, SC, or the following environmental perturbations on YPD medium: pH3, pH8, 0.01% Sodium dodecyl sulfate, 5 mM Caffeine (Caff), 6% Ethanol (EtOH), 100 mM CaCl_2_, 20 μg/ml Benomyl (Beno), 1.4 mM H_2_O_2_, 1.6 μg/ml Amphothericin B (Amph), 0.04% Methyl methanesulfonate (MMS), 80 mM Hydroxyurea, 150 μg/ml Fluconazole, 0.4 M KCl, 1 M Sorbitol, 1 M Glycerol, 50 μM Menadione, and 150 μM Paraquat dichloride. Reagents were purchased in Sigma Aldrich. These environments affect different cellular processes in yeast (Hampsey 1997; Dudley *et al.* 2005).

### Data analysis

Growth kinetics were analyzed as in (Warringer and Blomberg 2003) using Matlab. In brief, mean blank OD_600_ was subtracted to all OD_600_ values within the same plate; negative values were set to 10^−4^. Non-linearity at high OD_600_ values was corrected with the formula OD_corr_ = OD_600_ + 0.449(OD_600_)^2^ + 0.191(OD_600_)^3^. Smooth growth curves were obtained by eliminating all negative differential data points; smoothed OD_corr_ values were log10 transformed. A slope was calculated or each three consecutive time points in the growth curve. In a similar strategy than (Warringer and Blomberg 2003), doubling times (*D*) were calculated from the mean of the second and third maximum slopes of *log*_10_(*ODcorr*) as a function of time. Growth rate was defined as 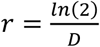. Mean growth rates 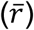 and its standard error from the independent replicates were calculated for each strain under each C_*j*_ cowth rate was defined ates (*G*) of KO_*i*_, and CM_*i*_ were defined as 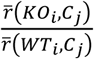 and 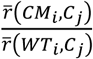, respectively.

For classification of catalytic and non-catalytic phenotypes, the growth rates of knockout (KO_*i*_) and catalytic-mutant (CM_*i*_) of each *GENE*_*i*_ under each C_*j*_ condition were tested for the null hypothesis 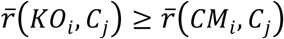 by a one-tailed Wilcoxon test (Matlab ranksum), with the alternative hypothesis 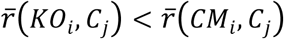. An FDR approach (Benjamini and Hochberg 1995) was used to correct *p*-values for multiple testing; "non-catalytic" phenotypes were defined using a 5% FDR threshold and values above this cutoff were classified as "catalytic". To avoid classification of marginal phenotypic effects, all cases where 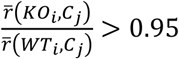 were labeled as "no phenotype".

The catalytic contribution to the phenotypes of each *GENE*_*i*_ under each condition *j* was defined as 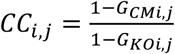, namely the ratio of the growth rates of catalytic-mutant and knockout strains relative to the WT_*i*_; *CC*_*i,j*_<0 (0.14% cases) and *CC*_*i,j*_ >1 (0.042% cases) were bound to 0 and 1, respectively.

### Differential epistasis profile analysis with genome-integrated mutants

We constructed the genomic-integrated variants of WT_*ILV1*_, KO_*ILV1*_, and CM_*ILV1*_. The *nat* cassette was amplified using A1 and A2 primers (**Table S4**) with sequences 160 bp downstream of *ILV1* and used to transform the parental Y8205 strain. DNA from the resulting strain (reference strain *ILV1*-*nat*) was used as a template for a PCR with primers ilv1-K109A and A2. The amplification product was transformed into Y8205 to obtain the catalytic mutant strain, *ilv1*^K109A^-*nat*. The knockout strain, *ilv1*∆::*nat*, was generated by replacing *ILV1* from the start codon to the same 160 bp downstream site using primers ILV1-MXF1 and A2.

Additionally, two control strains were constructed: an alternative knockout strain, *ilv1*∆(bis)::*nat*, obtained by replacement of *ILV1*’s ORF from the first to the last codon (with oligos ILV1-MXF1 and ILV1-MXF2), and another reference strain *NTR-nat* with a resistance marker integrated at a neutral locus, obtained by insertion of the *nat* cassette 200 bp downstream *ARO7* (oligos NTR-Nat-F and NTR-Nat-R). Strains with mutations integrated to the genome were used for the differential epistasis profile analysis (Bandyopadhyay *et al.* 2010; Braberg *et al.* 2013b). Reference, knockout, and catalytic mutant strains were mated to a collection of non-essential MATa *xxx*∆::*kan* deletion strains. Seven technical replicates for each reference strain *ILV1*-*nat* and *NTR*-*nat* were included, and one replicate of each *ilv1*^K109A^-*nat*, *ilv1*∆::*nat*, and *ilv1*∆(bis)::*nat* strain. After sporulation, each strain collection was triplicated, obtaining 21 of each reference-strain replicates, and three technical replicates for each of the three mutant constructs. Haploid MATa strains bearing the two resistance markers (*nat*, *kan*) were selected and pictures were taken after 48 hrs of incubation at 30°C in double-marker selection medium, as described in (Collins *et al.* 2006; Collins *et al.* 2010).

Genetic interaction scores (S-scores) were calculated from the sizes of the double-marker colonies as described, with the EMAP toolbox (Collins *et al.* 2006; Collins *et al.* 2010). S-scores account for the magnitude and the confidence of the genetic interaction. In brief, colony sizes were normalized by position and by the distribution of colony sizes in the plate. Given that genetic interactions are rare, the median of the normalized colony sizes of each plate and each array gene is used to calculate the expected double-marker colony size; the high number of technical replicates in reference strains *ILV1*-*nat* and *NTR*-*nat* were used to obtain robust data for such calculation (Schuldiner *et al.* 2005; Collins *et al.* 2010). Data from normalized colony sizes above or below an establish threshold were eliminated (size>1500; size<5). Also, data from double-marker strains was filtered out if 1) the replicates presented high standard errors, 2) if the mean S-score of either reference strain were above 2 or below ‒3, or 3) if the array gene was close to the *ILV1* locus (50 kb), avoiding apparent negative S-scores due to genetic linkage. The standard deviation of S-scores increases in a non-linear manner with respect to its magnitude (Bandyopadhyay *et al.* 2010); therefore, to compare the S-scores from different epistasis profiles, we calculated the variance (*σ*) as a non-parametric function using a sliding window of S-scores with the polynomial *σ* = 0.012x^2^ − 0.095x + 0.502, where *x* is de difference of the S-scores evaluated, with limits of *σ* at 2 (Bandyopadhyay *et al.* 2010).

### Functional clustering and GO-term enrichment

Functional modules were obtained for genes—from non-catalytic and catalytic genetic interactions separately—according to their shared functional terms (GO-terms and mutant-phenotype annotations, 1,748 terms in total). To do so, the overall agreement between gene-pairs was evaluated by Cohen’s kappa 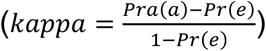, where *Pra*(*a*) is the relative observed agreement or the number of terms that a gene-pair shares divided by the total number of terms in the matrix, and *Pr*(*e*) is the probability of agreement by chance, calculated as the sum of probabilities for each member of the gene-pair to be associated or not to each term. Gene-pairs that showed *kappa* >0.35 were used as cluster seeds; groups sharing more than 50% of their genes were merged. The resulting gene modules were named based on the most common functional feature of genes in the module. Network representation was created using Cytoscape. GO-term annotations and mutant-phenotypes annotations were downloaded from the Yeast Genome Database (December 2016).

Gene Ontology enrichment analyses were performed using GOrilla (Eden *et al.* 2009), with a target list compared to the specific background list of genes tested. If specified, redundancy of GO-terms was removed using REVIGO (Supek et al. 2011).

All strains are available upon request. Table S1 contains a detailed description of all strains generated and used in this study. Table S3 and Table S4 describe all primers and their sequences. Phenotypic and genetic interaction data is provided in Dataset S1 and Dataset S2, respectively.

## RESULTS

### Selection of enzyme-coding genes and catalytic-mutant design

We set out to test in a systematic manner whether gene-deletion phenotypes of enzymes from *S. cerevisiae* are caused by the loss of their catalytic activities. To this end, we designed an experimental strategy based on the comparison of the growth phenotypes of catalytic-mutant strains (loss-of-function substitution of a single catalytic residue in an otherwise intact protein) and gene-deletion strains ('knockout', *i.e.* no protein at all) (**Figure 1**). We concentrated our experimental analysis on genes related to metabolism—a powerful model cellular system for studying gene function (Segre *et al.* 2005; Szappanos *et al.* 2011)—and focused our screen on enzymes of amino acid biosynthesis, since loss of their catalytic function can be readily confirmed by amino acid auxotrophy. From the Saccharomyces Genome Database (SGD, http://www.yeastgenome.org), we selected 86 single-gene knockouts (or double knockouts for paralogous isoenzymes) resulting in an amino acid auxotrophy; histidine auxotrophy was not considered because the parental strain used has a *his3*∆ genotype. Our experiments were based on haploid knockout strains complemented with centromeric plasmids; hence, out of the 86 auxotrophic and viable strains, we further considered 56 genes that were part of the MoBY-ORF plasmid collection with coding sequences from *S. cerevisiae* with their corresponding 5’-promoter and 3’-UTR sequences (Ho *et al.* 2009). Next, we selected enzymes annotated with a single molecular function (Ashburner *et al.* 2000) (the catalytic activity), and with well-characterized enzymatic reaction mechanisms in the Catalytic Site Atlas (Furnham *et al.* 2014) or the MACiE Database (Holliday *et al.* 2012). This resulted in 18 genes (herein referred to as the *GENE*_*i*_ set): *ALT1*, *ARG3*, *ARG4*, *ARO3*, *ARO4*, *ARO7*, *TKL1*, *TKL2*, *TRP1*, *ASN1*, *ASN2*, *ILV1*, *ILV2*, *BAT2*, *HOM2*, *HOM6*, *MET2*, and *THR4* (**Table 1**). Such enzymes perform a wide variety of reactions, represent diverse protein folds, and are involved in different amino acid biosynthetic pathways (**Table S2**).

**Figure 1.**
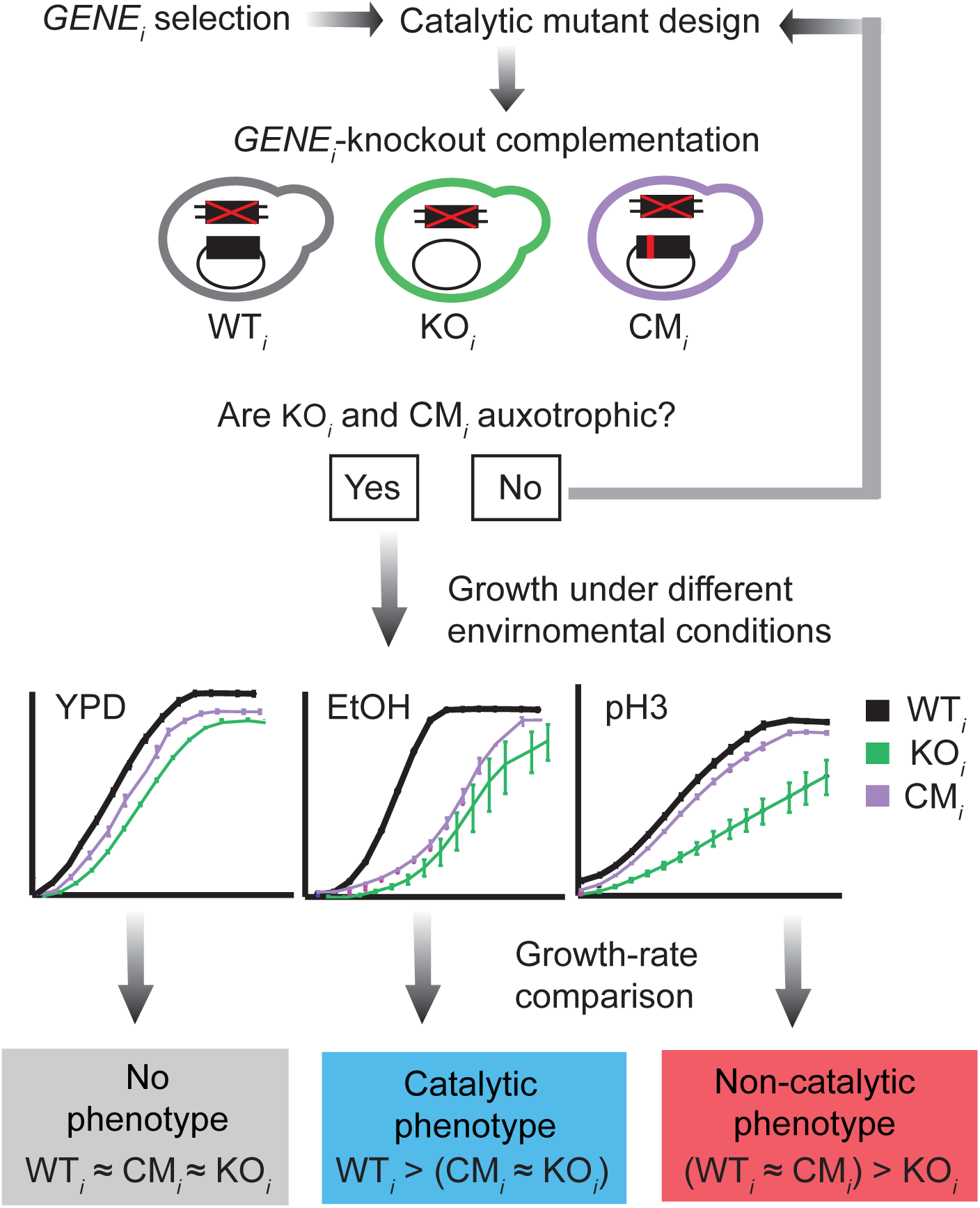
A systematic experimental strategy to dissect the molecular bases of enzyme-loss phenotypes. Enzyme-encoding genes (*GENE*_*i*_) from amino acid biosynthesis metabolism were selected; residues involved in early catalysis were targeted for site-directed mutagenesis. Plasmid-borne wild-type (WT_*i*_), gene knockout (KO_*i*_), or catalytic-mutant (CM_*i*_) constructs were used to complement the corresponding *gene*_*i*_Δ (deletion of *GENE*_*i*_). Loss of catalytic function was confirmed by amino acid auxotrophy; growth of complemented strains was characterized under different environmental conditions. Growth rates were used to classify each case as “no-phenotype” (gray), “catalytic phenotype” (cyan), or “non-catalytic phenotype” (red).

To generate strains expressing *GENE*_*i*_ proteins with no catalytic activity (catalytic mutant strain, CM_*i*_), we replaced a single “essential” catalytic residue that directly participates in the catalysis (see Materials and Methods). We tested the loss of catalytic activity by auxotrophy of *gene*_*i*_∆ strains bearing plasmids with site-directed mutations. Three out of the 18 *GENE*_*i*_ enzymes (Arg3, Hom2, Met2) were discarded because no site-directed mutant tested resulted in loss of catalysis. All other catalytic mutants were unable to grow after long incubation in minimal medium (**Figure 2A**; **Figure S1**). Although substitution of residues that are considered essential for enzymatic function may not completely abolish catalysis but rather alter the catalytic mechanism (Peracchi 2001), the fact that catalytic-mutant strains did not grow in the absence of amino acids indicates loss of the catalytic activity that is required for growth. Residual growth after 72 hrs of inoculation was only observed in the *ilv2*^E129A^ catalytic mutant. Moreover, many of the loss-of-function point mutations used here had been thoroughly characterized elsewhere (Fisher and Eisenstein 1993; Schnappauf *et al.* 1997; Delabarre *et al.* 2000; Kingsbury *et al.* 2015).

**Figure 2.**
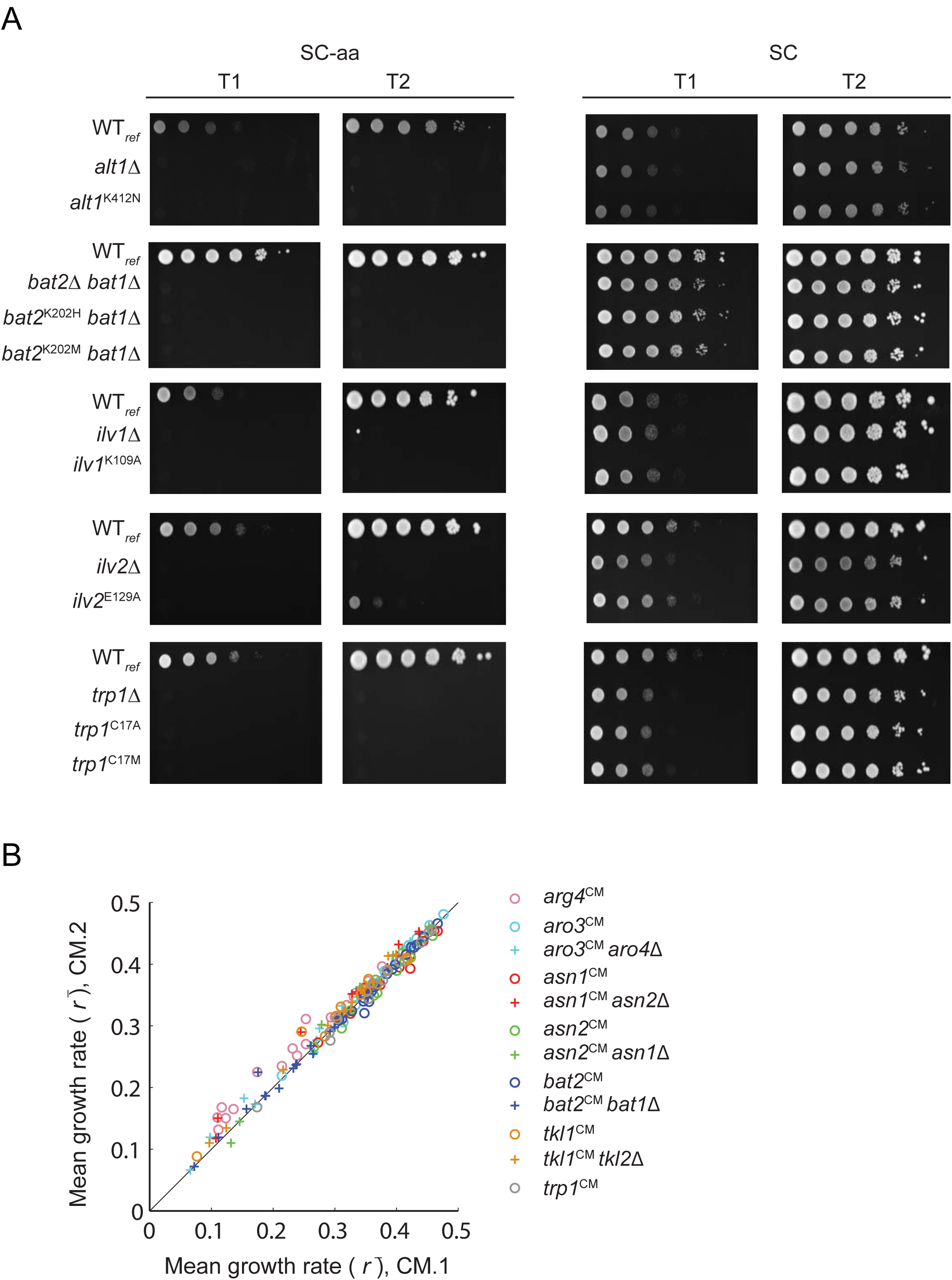
Complete loss of catalytic function with no additional dominant effects caused by specific residue replacements. **(A)**Drop-spot assays of auxotrophy. Culture dilutions were inoculated onto SC-aa (right) and SC-uracil (left) plates and incubated at 30°C. Images correspond to growth after 48 hrs (T1) and 96 hrs (T2) hrs for slow growers (*ALT1* and *BAT2*), or 24 hrs (T1) and 72 hrs (T2) for *ILV1*, *ILV2*, and *TRP2*. (**B**) Comparison of the mean growth rates under different environmental conditions (C_*j*_) of catalytic-mutant strains of *GENE*_*i*_ for which two different catalytic mutants were generated, validated, and screened (CM.1_*i*_ and CM.2_*i*_; *n*=165).

Site-directed substitutions of protein amino acids may impact organismal fitness by different mechanisms, in addition to the loss of a specific function (Tokuriki and Tawfik 2009; Jeffery 2011; Song *et al.* 2014). We therefore inspected if particular amino acid replacements in the catalytic mutants accounted for dominant effects on fitness. To this end, we compared the growth phenotypes of different amino acid substitutions of the same catalytic residue for enzymes in which two confirmed catalytic-mutant strains were available. Importantly, we observed high correlation of growth phenotypes between the two catalytic mutants characterized under different conditions (*r*^2^=0.98; **Figure 2B**). Such high correlation was not observed after randomizing mutant pairs (**Figure S2**). Taken together, these results suggest that the phenotypes of catalytic mutants are not the result of residual activity, altered catalytic properties, or dominant effects of particular amino acid replacements, but are more likely due to complete loss of enzyme catalysis.

### Quantitative analysis of the contribution of catalysis to phenotypes

For 15 *GENE*_*i*_ enzymes with a confirmed loss-of-function, auxotrophic CM_*i*_, we performed a large-scale phenotypic screen aimed to analyze the contribution of catalysis to the growth phenotypes of the corresponding gene knockout, KO_*i*_. To this end, we monitored the growth kinetics of all strains (**Table S1**) in five biological replicates (independent plasmid-transformation clones), challenged to 19 different growth conditions, C_*j*_ (see Materials and Methods). For each experiment, we calculated the growth rate and filtered out atypically high rates with respect to the WT reference (WT_ref_) grown in YPD (less than 0.4% samples; **Figure S3A**). As expected, growth under environmental perturbations was slower than on YPD (**Figure S3B**). To avoid hypomorphic effects associated with the expression of a gene from a centromeric construct, which would complicate downstream analysis, we filtered out genes in which the gene-specific WT_*i*_ strains grew slower than the WT_ref_ (4 out of 15 genes; **Figure S3C**). In the final data set of eleven *GENE*_*i*_ enzymes, growth of WT_*i*_ strains showed high correlation to the universal WT_ref_ strain (*r*^*2*^=0.905; **Figure S3D**). As expected, the median growth rate of KO_*i*_ and CM_*i*_ strains was significantly lower than growth of the WT_*i*_ (*p*<10^−15^ and *p*<10^−15^, respectively; one tailed Wilcoxon ranked-sum test). The complete phenotypic data set for the eleven *GENE*_*i*_ enzymes under 19 environmental conditions is provided in **Dataset S1**.

We classified the phenotypic data set in three groups based on the relative growth rates (*G*) of the KO_*i*_ and CM_*i*_ mutant strains: 1) no phenotype, 2) catalytic phenotype, and 3) non-catalytic phenotype. We found that in 38.6% out of 379 growth-rate comparisons, the knockout had little or no effect (*G*>0.95) (**Figure 3A**; "no phenotype"). We further classified the remaining 233 slow-growth phenotypes; for 70%, we found no significant difference between the growth phenotypes of catalytic and knockout mutants ("catalytic phenotypes"). Interestingly, in 30% cases the catalytic mutant grew significantly faster than the corresponding knockout (5% FDR, one-tailed Wilcoxon rank sum test). In such cases of "non-catalytic phenotypes" at least part of the phenotype did not depend on the loss of enzymatic activity. We note that catalytic-mutant strains rarely grew better than the wild type (1% *G*_CM*i*_>1.05) or worse than the corresponding knockout strain (6.8% *G*_CM*i*_ – *G*_KO*i*_<–0.05), which suggests that no structural defects or neomorphic phenotypes result from the site-directed mutations. Taken together, these results suggest that the loss of an additional molecular activity in the gene-knockout—a "moonlighting" function—could contribute to the observed phenotype.

**Figure 3.**
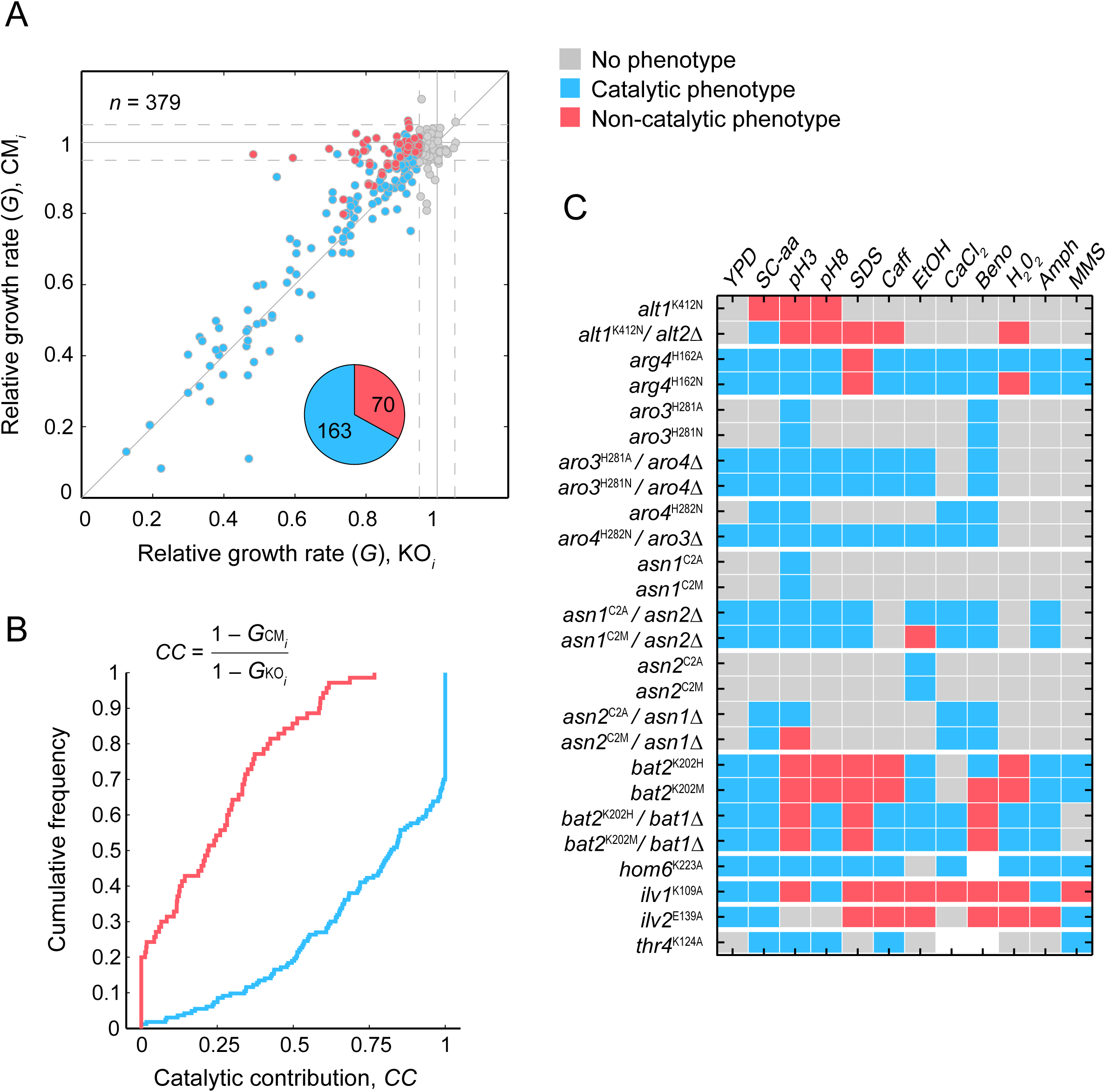
Non-catalytic phenotypes in biosynthetic enzymes from yeast. **(A)** Scatter plot of growth rates relative to WT_*i*_ (*G*) of gene knockouts KO_*i*_ (*x*-axis) and catalytic-mutants CM_*i*_ (*y*-axis) of eleven *GENE*_*i*_ characterized under different growth conditions. Growth phenotypes were defined as *G*<0.95 (gray dots indicate no phenotype); based on growth rate differences (5% FDR), phenotypes were classified as catalytic (cyan) and non-catalytic (red). Pie chart shows the fraction of catalytic and non-catalytic phenotypes. **(B)** Cumulative distribution of the catalytic contribution (*CC*) of the catalytic (cyan) and non-catalytic (red) phenotypes. **(C)** Figure shows the three phenotype categories of different *GENE*_*i*_ (vertical axis; labels are names of CM_*i*_ strains) under different environmental perturbations C_*j*_ (horizontal axis). White indicates missing data points. Only twelve conditions for which data was obtained for most of the *GENE*_*i*_ are shown.

As noted above, we defined non-catalytic phenotypes as significant slow growth of the KO_*i*_ compared to the CM_*i*_ strain. Therefore, in non-catalytic phenotypes at least part of the knockout phenotype is not explained by the loss of catalysis. To quantify the contribution of the loss of catalysis to the non-catalytic phenotypes, we established a catalytic contribution (*CC*) factor, result of the fraction of the magnitude of the knockout phenotype different to that of the catalytic mutant (see Materials and Methods). In this way, a *CC* value close to one means that catalysis solely explains the phenotype, while a value close to zero means that catalysis does not contribute to the gene-knockout phenotype. We observed that in the vast majority of the scored non-catalytic phenotypes the loss of the catalysis explained less than half of the knockout phenotype (*CC*=0.5) and in 20% of the cases catalysis did not contribute at all to the gene-deletion phenotype (*CC*=0; **Figure 3B**). As expected, the median catalytic contribution in scored catalytic phenotypes was high (*CC*=0.82; **Figure 3B**). These results underscore that, for an important number of observed phenotypes, the effect is driven majorly by the loss of a molecular function other than the known catalysis of the enzyme.

Most non-catalytic phenotypes were concentrated in deletions of *ALT1, BAT2, ILV1*, and *ILV2* (see **Figure S4** for KO_*i*_ and CM_*i*_ phenotypes). This result indicates that non-catalytic phenotypes are a feature of a limited set of genes encoding enzymes with possible moonlighting activities. Conversely, several genes showed consistent catalytic phenotypes (**Figure 3C**, cyan). Such was the case for knockouts of *ARO3, ARO4, ASN1, ASN2, HOM6*, and *THR4* (see **Figure S4** for examples). Catalytic profiles in which the sole loss of catalysis explains all or most growth phenotypes indicate a single molecular function, that is, a monofunctional protein, at least under the limited set of conditions tested.

Our screen included strains in which the corresponding duplicate-gene was deleted, which allowed us to test whether the presence of the paralog was masking the mutants' phenotypic effect. Indeed, duplicate genes *ALT1*, *ARO3*, *ARO4*, *ASN1, and ASN2* showed few growth phenotypes when analyzed in a single gene-knockout background. However, we observed more phenotypic diversity when the analysis was carried out in a double-knockout background (**Figure 3C**). For instance, the *aro3*∆ single-knockout had no phenotype under most conditions tested, but the double *aro3*∆*aro4*∆ strain grew slow under many conditions, which allowed us to describe the catalytic nature of the Aro3-depletion phenotype. Likewise, more phenotypes were scored in the *alt1*∆*alt2*∆ double knockout compared to the single *alt1*∆. Intriguingly, the exposed phenotypes were mostly non-catalytic; this is consistent with a lack of catalytic activity in the Alt2 paralog (Peñalosa-Ruiz *et al.* 2012). These results suggest that a moonlighting non-catalytic function is present in Alt1 and shared with its paralog. Overall, our phenotypic screen allowed us to identify a set of phenotypes that do not depend on catalysis, underscoring additional relevant functions of yeast enzymes.

### The non-catalytic genetic landscape of *ILV1*

To gain insight into the non-catalytic functions of one of the exposed moonlighting proteins, we focused on the genetic interactions of the *ILV1*-encoded threonine deaminase. This enzyme showed some of the strongest non-catalytic phenotypes in our screen. Measuring genetic interactions (epistasis), defined as the phenomenon in which the phenotype of a gene mutation is modified by the mutation of another gene, is a powerful way to reveal functional associations among genes (Segre *et al.* 2005; Boone *et al.* 2007; Costanzo *et al.* 2016). In particular, we generated a differential epistasis profile analysis (Bandyopadhyay *et al.* 2010; Braberg *et al.* 2013a) of knockout and catalytic variants of *ILV1* to describe the dependency on catalysis of its genetic interactions.

We generated genome-integrated constructs of wild-type (*ilv1*∆::*ILV1*-*NAT*), knockout (*ilv1*∆::*NAT*), and catalytic mutant (*ilv1*∆::*iv1*^K109A^-*NAT*). We mated each of these query strains to an array of 3,878 non-essential gene knockouts to finally obtain collections of double-mutant haploids (**Figure 4A**). Based on the colony sizes of the double-mutants compared to those of the corresponding single-knockout references, we obtained an S-score as a parameter of the magnitude and statistical confidence of each genetic interaction (Collins *et al.* 2006; Collins *et al.* 2010). Our epistasis profiles included an alternative reference strain with a different neutral-marker insertion site and an additional *ilv1*∆ strain with a different gene-deletion design (see Materials and Methods). Different query-strain libraries clustered as expected in terms of their S-score profiles (**Figure 4A**), while all mutant strains were similar in terms of colony-size variation within technical replicates (**Figure S5**). The distribution of S-scores of both mutant collections (*ilv1*∆ and *ilv1*^K109A^) was centered at zero, with a short tail of positive (alleviating) and a long tail of negative (aggravating) genetic interactions (**Figure 4B**). Genetic interactions were defined using a fixed cutoff of |S-score|>3 (Bandyopadhyay *et al.* 2010) in either double mutant collection. Using this criterion, we found that in the combination of both mutant collections, 9% double-mutants resulted in negative interactions, while positive interactions were scored in 2% of the cases (**Figure 4B**; **Dataset S2**). These genetic interactions were enriched primarily in genes of metabolism, mitochondrial function, and chromatin organization (**Table S5**).

**Figure 4.**
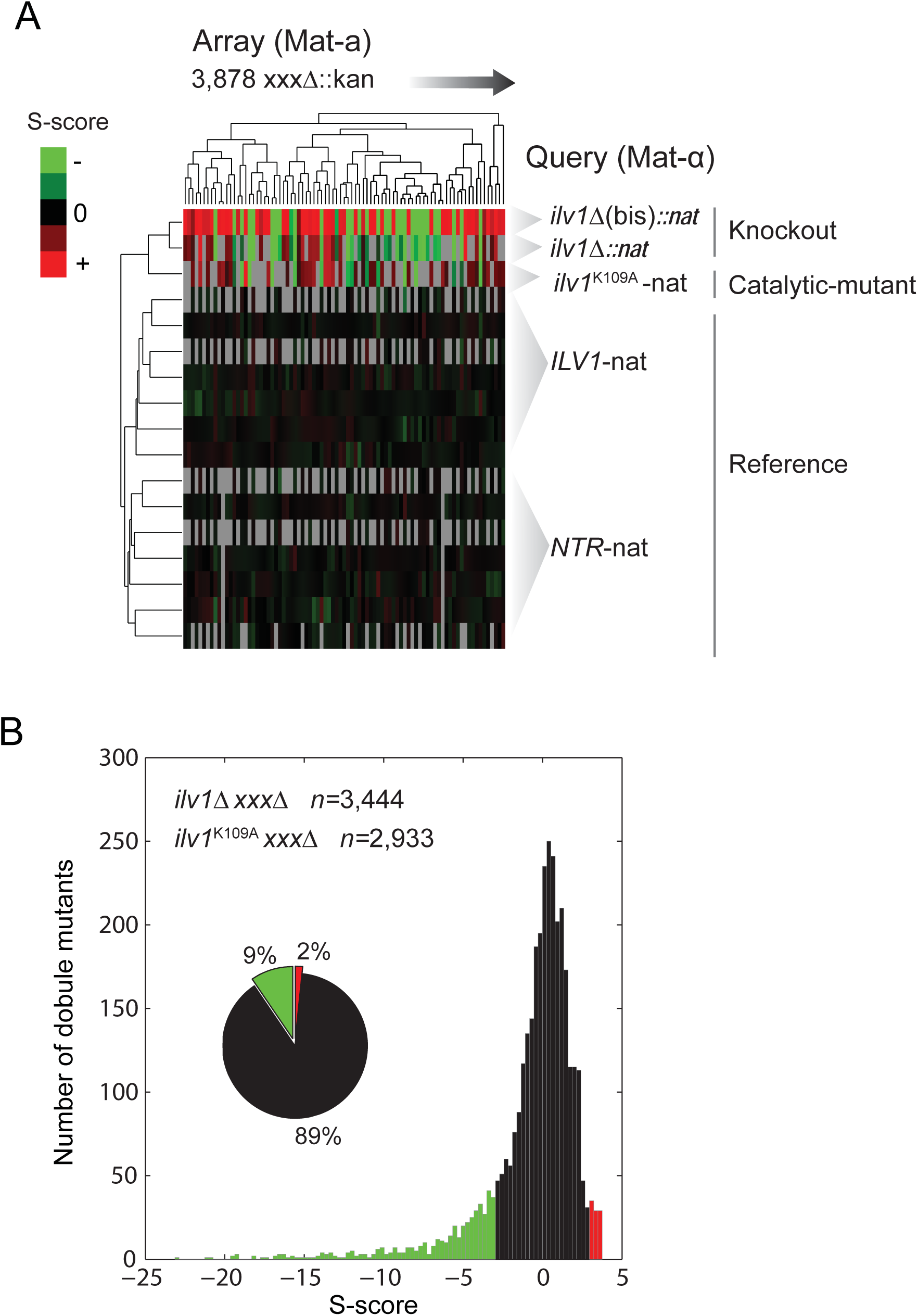
Epistasis-profile data analysis of *ILV1*. **(A)** Double-mutant and reference collections were generated by mating Mat-α query strains of genome-integrated variants of *ILV1* with an array of Mat-a single-knockouts of most non-essential genes (see Materials and Methods). Heat-map shows hierarchical bidimensional clustering of array genes (*x*-axis) and query genes (*y*-axis) by their S-score profiles. Clustering was performed by average linkage with a Spearman-rank similarity metric using Gene Cluster 3.0; only a small subset of array genes is shown. **(B)** Histogram of S-scores from filtered data of the knockout and catalytic-mutant strain collections (*ilv1*∆ *xxx*∆ and *ilv1*^K109A^ *xxx*∆). Negative (green) and positive (red) genetic interactions in either collection are shown, as defined with a cutoff of |S-score|>3.

To describe the catalytic dependency of the genetic-interaction landscape of *ILV1*, we compared the S-score profiles of the *ilv1*∆ and *ilv1*^K109A^ collections (**Figure 5A**). Strikingly, we observed a wide dispersion of S-scores above the diagonal, indicating that the strength of some negative genetic interactions in the *ilv1*∆ knockout was diminished in the *ilv1*^K109A^ catalytic mutant. This trend was not observed when contrasting the S-scores of the two different knockout strains (*ilv1*∆ and *ilv1*∆(bis); **Fig S6A**). These results suggested that an important number of genetic interactions of *ILV1* do not depend on loss of its catalytic activity.

**Figure 5.**
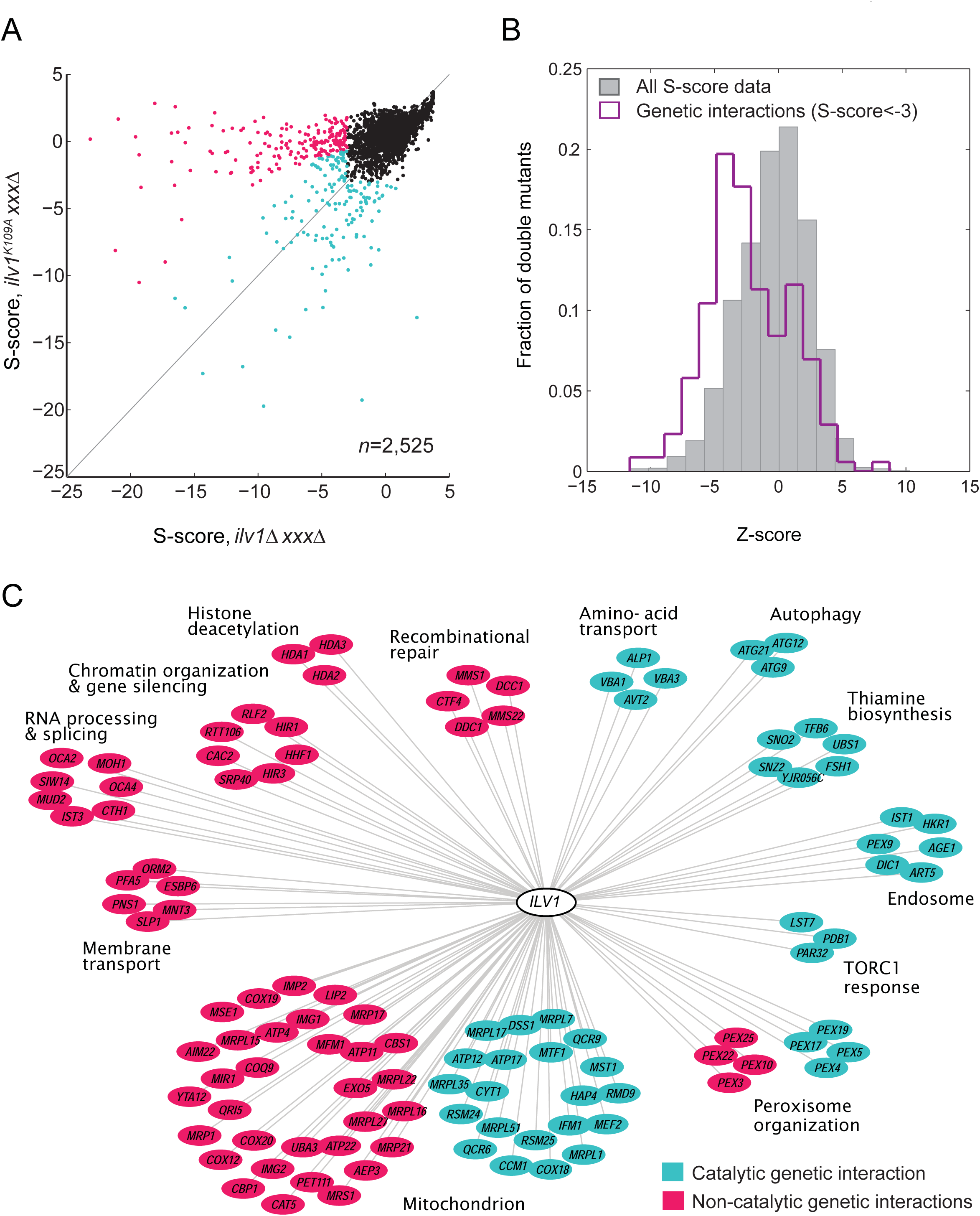
A set of genetic interactions of *ILV1* are not driven by loss of catalysis. **(A)** A differential epistasis-profile analysis of knockout and catalytic mutants of *ILV1*. Scatter plots of S-scores for the *ilv1*∆ knockout (horizontal axis) and the *ilv1*^K109A^ catalytic-mutant (vertical axis) collections. Negative genetic interactions were defined by S-score<‒3 in either collection. Non-catalytic genetic interactions (magenta) were scored based on a stronger negative S-score of the knockout (*p*<0.005). Negative genetic interactions with no significant difference (catalytic interactions) are shown in cyan. **(B)** Histogram of *Z*-scores of all compared gene pairs (gray bars) and gene pairs with significant negative genetic interactions (purple line). **(C)** Network representation of functional modules (*kappa*>0.35) of genes with non-catalytic (magenta) or catalytic (cyan) genetic interactions with *ILV1*.

To identify the specific cases of non-catalytic genetic interactions of *ILV1*, we performed a differential epistasis-profile analysis by defining a *Z*-score of the difference in the corresponding S-scores (**Figure S6B**). We focused only on negative interactions, to avoid over-scoring of marginal differences that could result from the narrow dynamic range of the positive genetic-interaction spectrum. The *Z*-scores of genes with negative genetic interactions were skewed to negative values (**Figure 5B**). Indeed, we found 187 (54.2%) non-catalytic interactions out of 345 negative gene interactions (*p*<0.005); the remaining 158 were defined as catalytic interactions.

Non-catalytic genetic interactions could arise from atypical features of the *ilv1*^K109A^ collection leading to a bias towards positive S-scores. We therefore inspected the colony sizes and their coefficients of variation in the double mutants (**Figure S6C**), which indicated that non-catalytic genetic interactions did not depend on unusual large colonies (leading to positive S-scores) or unusual high standard deviations (resulting in S-scores close to zero). In sum, these observations reveal that around one half of the genetic interactions of *ILV1* do not depend solely on its catalytic activity, suggesting that, indeed, such non-catalytic interactions are driven by the phenotypes arising from the loss of moonlighting activities of *ILV1*.

To describe the cellular functions associated to both interaction categories, namely catalytic and non-catalytic genetic interactions, we grouped genes from each category separately according to their shared GO-terms and mutant-phenotype annotations (*kappa*<0.35; **Figure 5C**). Both catalytic and non-catalytic hits were grouped in modules of genes with mitochondrial and peroxisome function, while catalytic hits resulted in clusters of genes involved in amino acid transport, autophagy, and TOR1-mediated response. Interestingly, many non-catalytic hits were clustered in different modules of genes with no direct connection to amino acid metabolism, RNA processing and splicing, chromatin organization, gene silencing, and the HDA1 complex. Such functional non-catalytic connections of *ILV1* to chromatin modification were also observed by GO-term enrichment analysis (**Table S6**). The genetic interactions of the *ILV1*-deletion with genes of chromatin regulation had been scored before in genome-wide epistasis screens (Costanzo *et al.* 2016). Remarkably, the genetic-interaction profiles of *ILV1* with chromatin remodelers is similar to those of genes involved in stress response (*RIM13, RIM20, WHI2*), protein sorting (*SPR3, CHS5, SEC63*), and chromatin remodeling (*ISW2, POB3*), but not to those of other metabolic genes (Bellay *et al.* 2011). Taken together, these observations indicate that the non-catalytic moonlighting activity of *ILV1* is associated to gene regulation, specifically to chromatin modification, in response to stress and other stimuli.

## DISCUSSION

Molecular biology has undergone a paradigm shift from the one gene – one function paradigm, while genetics and genomics have provided systems views of genes and proteins in their cellular context. Even though these advancements have led to the awareness of different types of gene multifunctionality, we are still in the quest to identify and reveal their underlying molecular and cellular mechanisms. Here, we focused on amino acid biosynthesis metabolism as a model system and dissected the gene-deletion phenotypes of enzymes into those that can be explained by the loss of catalysis and those that cannot. We screened the phenotypic profiles of *S. cerevisiae* gene-deletion, catalytic-mutant, and reference strains, and found that as many as 30% of the gene-deletion phenotypes tested were non-catalytic. Such cases, in which loss of the catalytic function did not recapitulate loss of the corresponding enzyme, suggest proteins with a “moonlighting” behavior, and were prevalent in four enzymes: Alt1, Bat2, Ilv1, and Ilv2.

Our finding that most gene-deletion phenotypes tested were driven by their catalytic function is in agreement with the view that genetic pleiotropy is usually caused by the perturbation of a single molecular function that affects many different cellular traits (He and Zhang 2006). Nonetheless, identifying non-catalytic functions could be challenging in conditions of strong dependence on the catalytic function. For example, the branched-chain amino acid transaminase, Bat2, showed non-catalytic phenotypes in the single-gene knockout, that were rendered strongly catalytic in the double *bat1*Δ *bat2*Δ background. Moreover, mutations in *HOM6* and *THR4* are known to be highly pleiotropic because of the accumulation of toxic metabolic intermediates (Arévalo-Rodríguez *et al.* 2004; Kingsbury and McCusker 2010). In such cases, additional mutations in the metabolic pathway would provide a better means to interrogate the extent to which pleiotropy is explained solely by catalysis. Genes with consistent catalytic profiles in our screen should therefore be considered monofunctional until proven otherwise.

Four out of eleven amino acid biosynthesis enzymes tested showed a moonlighting behavior. Previous studies have suggested that the moonlighting hits in our screen could indeed have more than one molecular function. For instance, early studies had proposed that the *ILV1*-encoded enzyme from yeast is a multifunctional protein involved both in catalysis and in the regulation of the expression of genes of isoleucine and valine biosynthesis (Bollon and Magee 1971; Calhoun 1976). In addition, Bat1 and Bat2 control TORC1 signaling through a non-catalytic structural function (Kingsbury *et al.* 2015). Interestingly, the branched-chain aminotransferases have been retained throughout the evolution of metazoans even though the anabolic pathways in which it participates have been lost—this is also the case for Ilv2, also identified as a moonlighting in our screen (Costa *et al.* 2015). We also note that the alanine transaminase Alt1 is a regulator of yeast chronological lifespan through metabolic-flux control (Yu *et al.* 2013), but whether catalysis is enough for this biological role has not yet been directly addressed. The quantitative nature of our genetic screen allowed us to identify some cases in which catalysis contributed partially to a “non-catalytic” phenotype. Moonlighting proteins are usually defined as molecules with two or more activities that are independent from one another (Huberts and van der Klei 2010; Zanzoni *et al.* 2015; Khan and Kihara 2016); however, our observations suggest that such activities may sometimes not be completely uncoupled. Partial functional contribution to phenotypes would be expected if two molecular activities in one protein crosstalk to each other in the broader cellular context, for example in the case of enzymes that moonlight by acting as direct transcriptional regulators of genes in the same metabolic pathway (Meyer *et al.* 1991; Moore *et al.* 2003). Alternatively, single site-directed mutations affecting two molecular activities in moonlighting proteins could result in partial phenotypic contributions. The quantitative genetic description of moonlighting proteins will provide further understandings of how multiple activities originate, coexist, and evolve in a single polypeptide.

In conclusion, our study shows that the gene-loss phenotypes of metabolic enzymes frequently do not depend on an annotated catalytic activity. We have learned that the chance to uncover proteins with such moonlighting behavior is relative to the magnitude, frequency, and regularity of the phenotype under different cellular contexts and in our ability to detect and measure it. These characteristics, in turn, depend on genetic redundancy, the degree of molecular and phenotypic interdependence between functions in a single polypeptide, and the dominance of the phenotypes associated to multifunctionality. In addition, genetic-interaction screens of site-directed mutants in yeast provide a powerful means to uncover mechanisms of protein multifunctionality, shedding light on the cellular roles of unknown moonlighting functions. Even though we focused on enzymes, our strategy can be readily used to identify other types of potentially-multifunctional proteins. Most likely, numberless moonlighting proteins within and beyond metabolism are yet to be discovered and characterized, providing a deeper understanding of cell biology, from metabolism and functional annotation to single gene and complex traits.

## Acknowledgements

We thank help from C. Abreu-Goodger with statistical analyses and F. Barona-Gómez with catalytic-mutant design. We are grateful to C. Abreu-Goodger and E. Mancera for critical reading of the manuscript. Strain BUN20 and plasmid p5586 were kindly shared by Charles Boone's laboratory. This work was funded by the Consejo Nacional de Ciencia y Tecnología de México (CONACYT grant CB2015/164889 to A.D.) and the University of California Institute for Mexico and the United States (UC MEXUS-COANCYT grant CN15-48 to A.D.). A.E.-C. had a doctoral fellowship from CONACYT. The funders had no role in study design, data collection and analysis, decision to publish, or preparation of the manuscript.

## SUPPLEMENTAL MATERIAL

**Figure S1.**
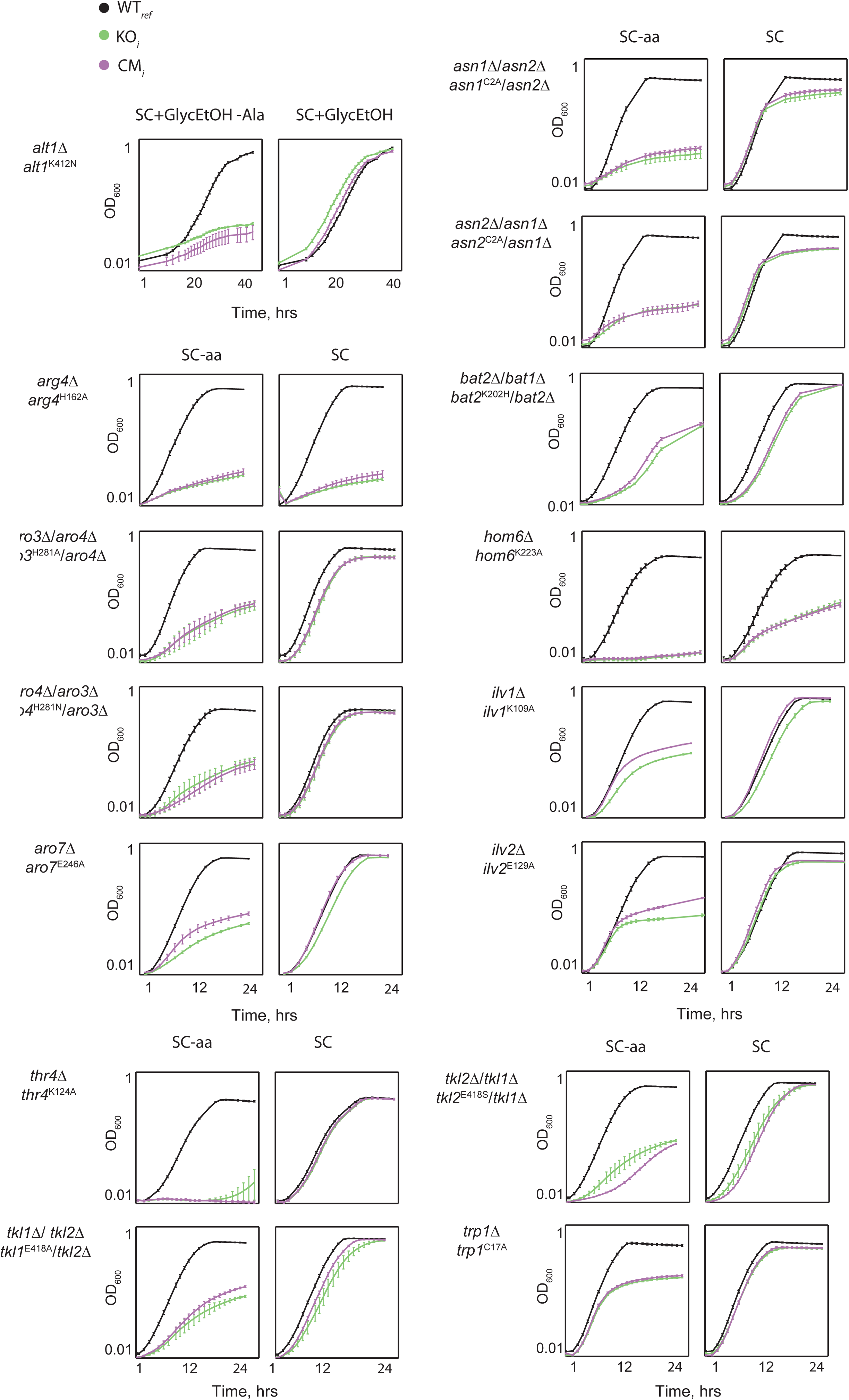
Confirmation of catalytic-inactive mutants by amino acid auxotrophy. Each pair of plots show growth curves of WT_*ref*_ (black) *KO*_*i*_ (green), and CM_*i*_ (magenta) strains under SC-aa (specific auxotrophy test) or SC medium. The specific amino acid dropout used in each case is indicated in Table 1. Data points are the average of five experimental replicates (different colonies from plasmid complementation); error bars are the S.E.M. Only one catalytic mutant is shown for each *GENE*_*i*_. *ALT1* auxotrophy was tested in SC medium with 3% glycerol and 2% ethanol without alanine; the *alt1*∆ single knockout was used given that the *ALT2* paralog has no alanine transaminase activity (Peñalosa-Ruiz *et al.* 2012). Drop-spot assays of auxotrophy were carried out for cases in which either the CM_*i*_ or KO_*i*_ strain grew at a final OD_600_>0.3 under SC-aa (see **Figure 1A**).

**Figure S2.**
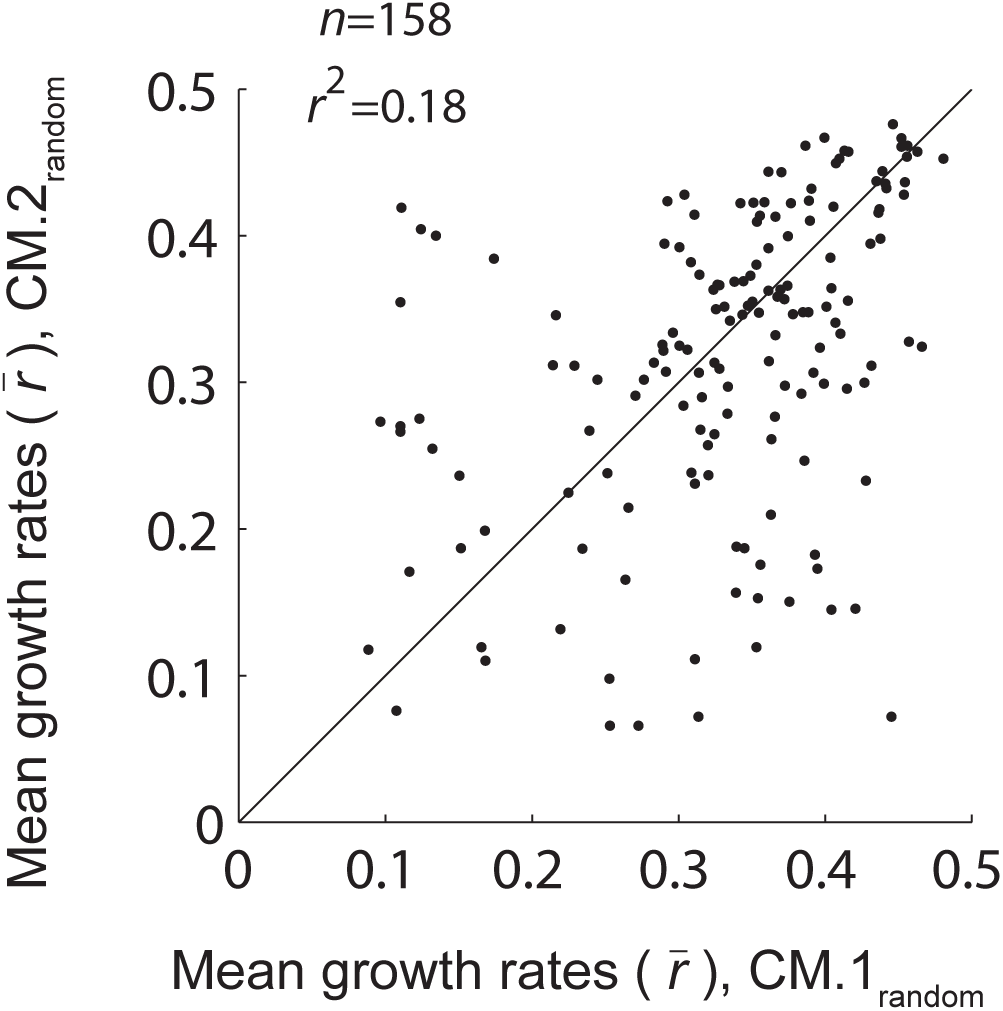
Random association of catalytic mutants. Scatter plot of mean growth rates of randomized pairs of catalytic mutants under fixed (non randomized) environmental conditions. Random pairs of catalytic mutants were obtained using Matlab (randperm) from the seven *GENE*_*i*_ for which two different catalytic mutants were generated.

**Figure S3.**
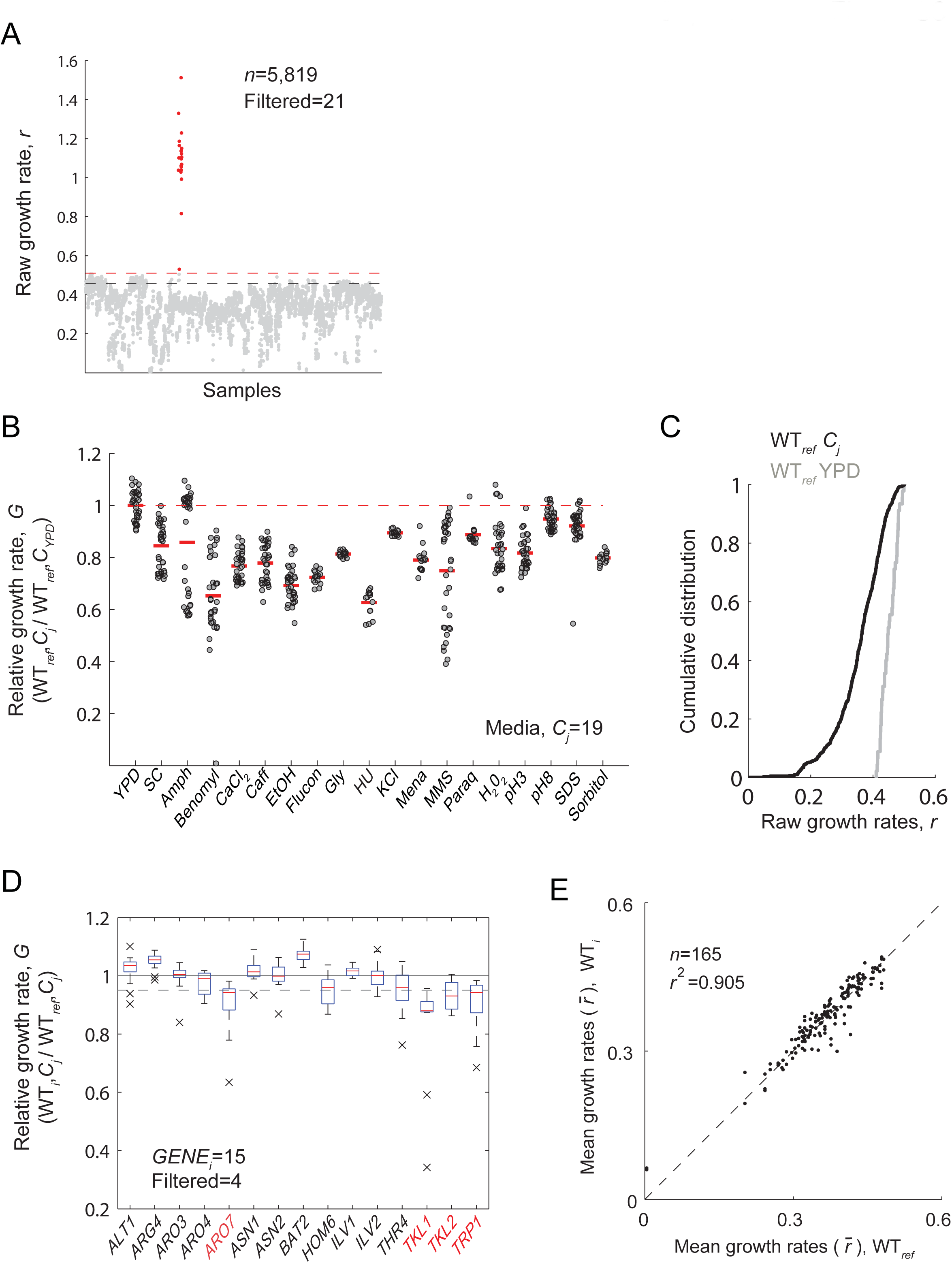
Data processing and quality control. **(A)** Raw growth rates of all samples. An upper-bound was defined as 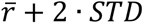 from 53 WT_ref_ strains grown in YPD. All *r*_*ij*_ data beyond this upper bound were eliminated (red dots). **(B)** Relative growth rates of the WT_ref_ strain under different environmental conditions, C_*j*_; solid red lines indicate median values. The bimodality of samples in some conditions is due to batch effects; comparisons of WT_*i*_, KO_*i*_, and CM_*i*_ were always done with strains from the same experiment batch. **(C)** Cumulative distributions of WT_*ref*_ raw growth rates in different environmental perturbations C_*j*_ (black line) and in YPD (grey line). **(D)** Plot shows growth rates of the references for each *GENE*_*i*_ (WT_*i*_ strains) under different conditions. A *GENE*_*i*_ was not further considered if the median of WT_*i*_ average growth rates 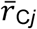 was too low compared to WT_*ref*_ growth: 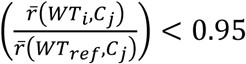. Genes *ARO7*, *THR1*, *TKL2*, and *TRP1* (red) were filtered out of the analysis. **(E)** Scatter plot of average growth rates of WT_*ref*_ and WT_*i*_ (final processed data).

**Figure S4.**
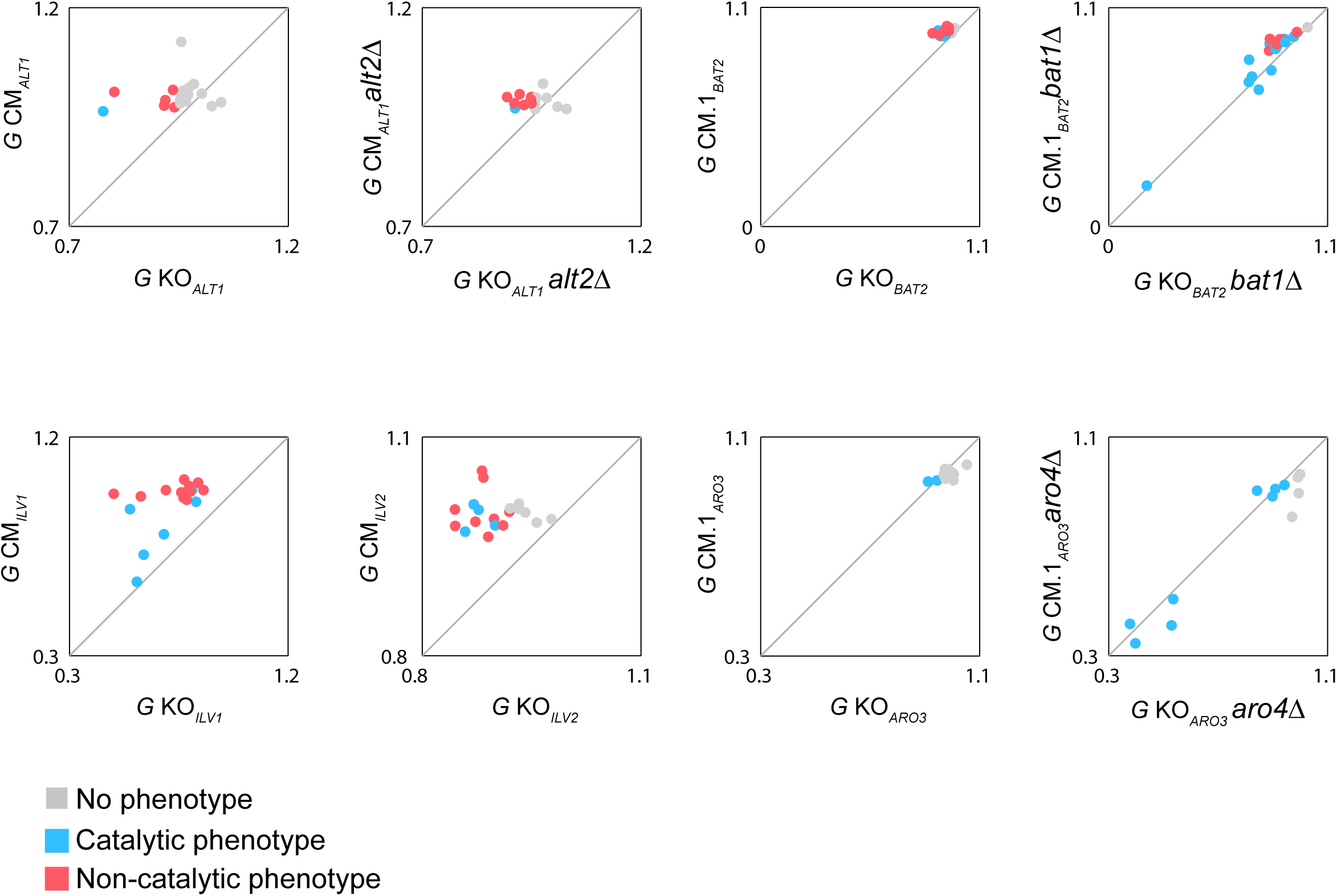
Scatter plots of gene-deletion and catalytic mutants for selected *GENE*_*i*_. The relative growth rates (*G*) of gene-knockouts (KO_*i*_, *x*-axis) and catalytic-mutants (CM_*i*_, *y*-axis) under different growth conditions. Colors indicate the phenotypic classifications: No phenotype (gray), catalytic phenotype (cyan), or non-catalytic phenotype (magenta).

**Figure S5.**
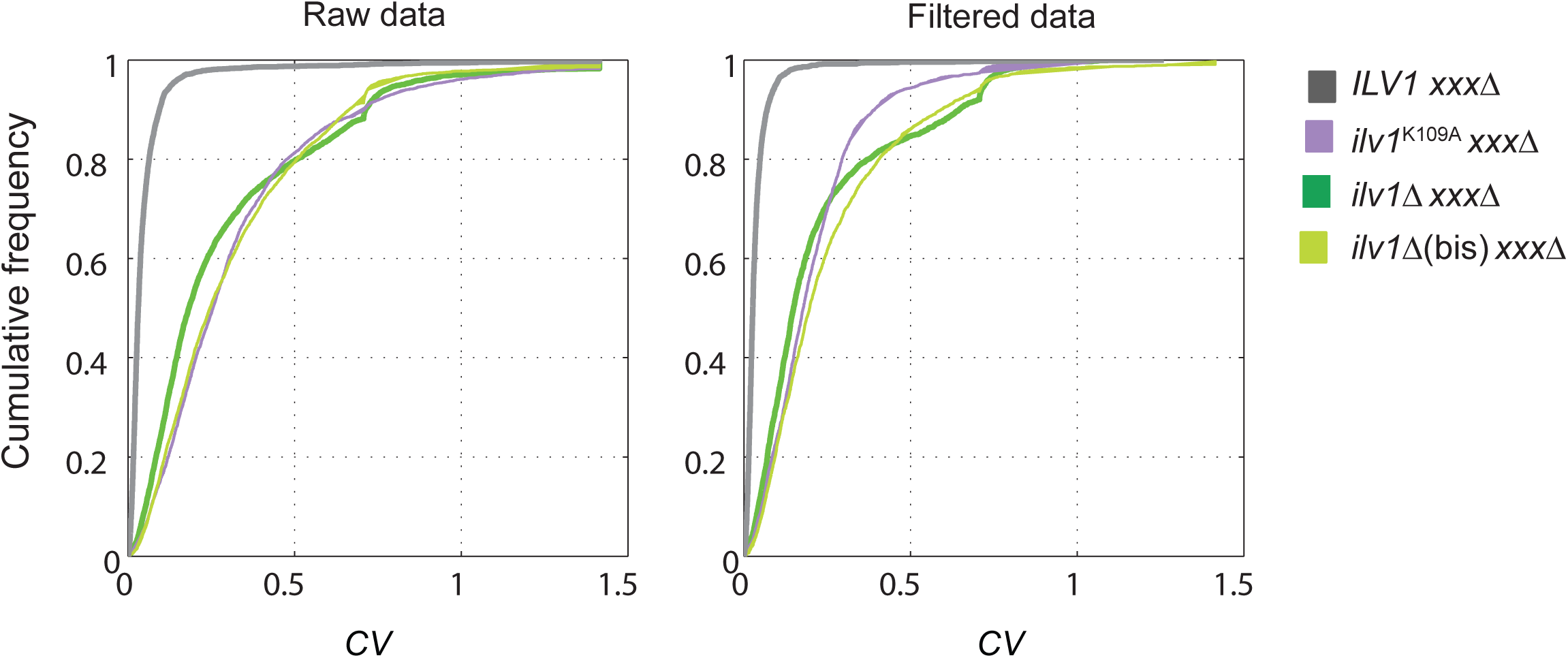
Colony size variation in epistasis-profile data. Cumulative distribution of the coefficients of variation (*CV*) of colony sizes in three technical replicates from raw (left) or filtered data (right). The distribution of *CV* between the double mutants from knockouts (*ilv1*∆ *xxx*∆ and *ilv1*∆(bis) *xxx*∆) and catalytic-mutant (*ilv1*^K109A^ *xxx*∆) strain collections were similar between each other before and after filtering the data; no bias in S-scores is expected from the variance in different epistasis profiles.

**Figure S6.**
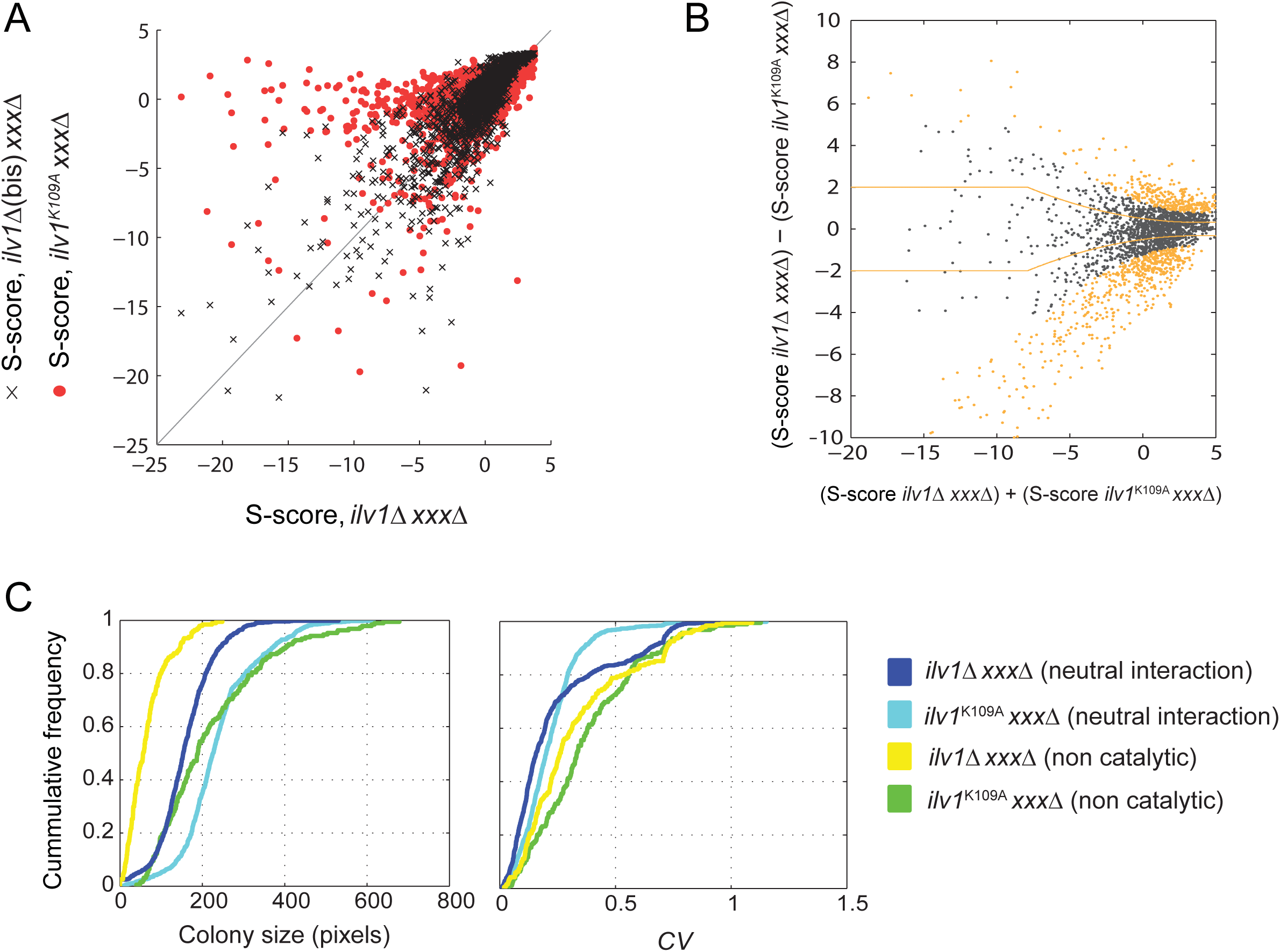
Differential epistasis-profile analysis. **(A)** Scatter plots of S-scores of the collections of knockout strain *ilv1*∆ and the catalytic-mutant *ilv1*^K109A^ (red dots) or the alternative knockout strain *ilv1*∆(bis) *xxx*∆ (black crosses). **(B)** The difference in S-scores of the *ilv1*∆ and *ilv1*^K109A^ collections is plotted as a function of their sum. A sliding window was used to calculate the global variance (orange lines) as described (Bandyopadhyay *et al.* 2010), which is used to obtain a *Z*-score for each (S-score *ilv1*∆)‒(S-score *ilv1*^K109A^) difference. Significant differences are highlighted (*p*<0.005; orange dots). **(C)** Distributions of colony size (left) and its coefficient of variation (right) of the knockout (*ilv1*∆) and catalytic (*ilv1*^K109A^) mutant collections for neutral interactions (|S-score|<1) or non-catalytic genetic interactions (S-score<‒3, *p*<0.005).

**Table S1.** Strains generated and used in this study.

| *GENE<sub>i</sub>* | Genotype <sup>a</sup> | Plasmid insert <sup>b</sup> | Strain |
| --- | --- | --- | --- |
| - | *his3Δ::nat<sup>R</sup>* | Φ | WT<sub>ref</sub> |
| *ALT1* | *alt1Δ::nat<sup>R</sup>* | *ALT1* | WT<sub>ALT1</sub> |
|  |  | Φ | KO<sub>ALT1</sub> |
|  |  | *alt1<sup>K412N</sup>* | CM<sub>ALT1</sub> |
|  | *alt1Δ::nat<sup>R</sup> alt2Δ::hyg<sup>R</sup>* | *ALT1* | WT<sub>ALT1</sub> *alt2Δ* |
|  |  | Φ | KO<sub>ALT1</sub> *alt2Δ* |
|  |  | *alt1<sup>K412N</sup>* | CM<sub>ALT1</sub> *alt2Δ* |
| *ARG4* | *arg4Δ::nat<sup>R</sup>* | *ARG4* | WT<sub>ARG4</sub> |
|  |  | Φ | KO<sub>ARG4</sub> |
|  |  | *arg4<sup>H162A</sup>* | CM.1<sub>ARG4</sub> |
|  |  | *arg4<sup>H162N</sup>* | CM.2<sub>ARG4</sub> |
| *ARO3* | *aro3Δ::nat<sup>R</sup>* | *ARO3* | WT<sub>ARO3</sub> |
|  |  | Φ | KO<sub>ARO3</sub> |
|  |  | *aro3<sup>H281A</sup>* | CM.1<sub>ARO3</sub> |
|  |  | *aro3<sup>H281N</sup>* | CM.2<sub>ARO3</sub> |
|  | *aro3Δ::nat<sup>R</sup> aro4Δ::hyg<sup>R</sup>* | *ARO3* | WT<sub>ARO3</sub> *aro4Δ* |
|  |  | Φ | KO<sub>ARO3</sub> *aro4Δ* |
|  |  | *aro3<sup>H281A</sup>* | CM.1<sub>ARO3</sub> *aro4Δ* |
|  |  | *aro3<sup>H281N</sup>* | CM.2<sub>ARO3</sub> *aro4Δ* |
| *ARO4* | *aro4Δ::nat<sup>R</sup>* | *ARO4* | WT<sub>ARO4</sub> |
|  |  | Φ | KO<sub>ARO4</sub> |
|  |  | *aro4<sup>H282N</sup>* | CM<sub>ARO4</sub> |
|  | *aro3Δ::nat<sup>R</sup> aro4Δ::hyg<sup>R</sup>* | *ARO4* | WT<sub>ARO4</sub> *aro3Δ* |
|  |  | Φ | KO<sub>ARO4</sub> *aro3Δ* |
|  |  | *aro4<sup>H282N</sup>* | CM<sub>ARO4</sub> *aro3Δ* |
| *ARO7* | *aro7Δ::nat<sup>R</sup>* | *ARO7* | WT<sub>ARO7</sub> |
|  |  | Φ | KO<sub>ARO7</sub> |
|  |  | *aro7<sup>E246A</sup>* | CM<sub>ARO7</sub> |
| *ASN1* | *asn1Δ::nat<sup>R</sup>* | *ASN1* | WT<sub>ASN1</sub> |
|  |  | Φ | KO<sub>ASN1</sub> |
|  |  | *asn1<sup>C2A</sup>* | CM<sub>ASN1</sub> |
|  |  | *asn1<sup>C2M</sup>* | CM<sub>ASN1</sub> |
|  | *asn1Δ::nat<sup>R</sup> asn2Δ::hyg<sup>R</sup>* | *ASN1* | WT<sub>ASN1</sub> *asn2Δ* |
|  |  | Φ | KO<sub>ASN1</sub> *asn2Δ* |
|  |  | *asn1<sup>C2A</sup>* | CM.1<sub>ASN1</sub> *asn2Δ* |
|  |  | *asn1<sup>C2M</sup>* | CM.2<sub>ASN1</sub> *asn2Δ* |
| *ASN2* | *asn2Δ::nat<sup>R</sup>* | *ASN2* | WT<sub>ASN2</sub> |
|  |  | Φ | KO<sub>ASN2</sub> |
|  |  | *asn2<sup>C2A</sup>* | CM.1<sub>ASN2</sub> |
|  |  | *asn1<sup>C2M</sup>* | CM.2<sub>ASN2</sub> |
|  | *asn1Δ::nat<sup>R</sup> asn2Δ::hyg<sup>R</sup>* | *ASN2* | WT<sub>ASN2</sub> *asn1Δ* |
|  |  | Φ | KO<sub>ASN2</sub> *asn1Δ* |
|  |  | *asn2*<sup>C2A</sup><br>*asn2*<sup>C2M</sup> | CM.1<sub>ASN2</sub> *asn1*Δ<br>CM.2<sub>ASN2</sub> *asn1*Δ |
| *BAT2* | *bat2*Δ::nat<sup>R</sup> | *BAT2*<br>Φ<br>*bat2*<sup>K202H</sup><br>*bat2*<sup>K202M</sup> | WT<sub>BAT2</sub><br>KO<sub>BAT2</sub><br>CM.1<sub>BAT2</sub><br>CM.2<sub>BAT2</sub> |
|  | *bat2*Δ::nat<sup>R</sup> *bat1*Δ::hyg<sup>R</sup> | *BAT2*<br>Φ<br>*bat2*<sup>K202H</sup><br>*bat2*<sup>K202M</sup> | WT<sub>BAT2</sub> *bat1*Δ<br>KO<sub>BAT2</sub> *bat1*Δ<br>CM.1<sub>BAT2</sub> *bat1*Δ<br>CM.2<sub>BAT2</sub> *bat1*Δ |
| *HOM6* | *hom6*Δ::nat<sup>R</sup> | *HOM6*<br>Φ<br>*hom6*<sup>K223V</sup> | WT<sub>HOM6</sub><br>KO<sub>HOM6</sub><br>CM<sub>HOM6</sub> |
| *ILV1* | *ilv1*Δ::nat<sup>R</sup> | p*ILV1*<br>Φ<br>*ilv1*<sup>K109A</sup> | WT<sub>ILV1</sub><br>KO<sub>ILV1</sub><br>CM<sub>ILV1</sub> |
| *ILV2* | *ilv2*Δ::nat<sup>R</sup> | *ILV2*<br>Φ<br>*ilv2*<sup>E139A</sup> | WT<sub>ILV2</sub><br>KO<sub>ILV2</sub><br>CM<sub>ILV2</sub> |
| *THR4* | *thr4*Δ::nat<sup>R</sup> | *THR4*<br>Φ<br>*thr4*<sup>K124A</sup> | WT<sub>THR4</sub><br>KO<sub>THR4</sub><br>CM<sub>THR4</sub> |
| *TKL1* | *tkl1*Δ::nat<sup>R</sup> | *TKL1*<br>Φ<br>*tkl1*<sup>E418A</sup><br>*tkl1*<sup>E418S</sup> | WT<sub>TKL1</sub><br>KO<sub>TKL1</sub><br>CM.1<sub>TKL1</sub><br>CM.2<sub>TKL1</sub> |
|  | *tkl1*Δ::nat<sup>R</sup> *tkl2*Δ::hyg<sup>R</sup> | *TKL1*<br>Φ<br>*tkl1*<sup>E418A</sup><br>*tkl1*<sup>E418S</sup> | WT<sub>TKL1</sub> *tkl2*Δ<br>KO<sub>TKL1</sub> *tkl2*Δ<br>CM.1<sub>TKL1</sub> *tkl2*Δ<br>CM.2<sub>TKL1</sub> *tkl2*Δ |
| *TKL2* | *tkl2*Δ::nat<sup>R</sup> | *TKL2*<br>Φ<br>*tkl2*<sup>E418S</sup> | WT<sub>TKL2</sub><br>KO<sub>TKL2</sub><br>CM<sub>TKL2</sub> |
|  | *tkl1*Δ::nat<sup>R</sup> *tkl2*Δ::hyg<sup>R</sup> | *TKL2*<br>Φ<br>*tkl2*<sup>E418S</sup> | WT<sub>TKL2</sub> *tkl1*Δ<br>KO<sub>TKL2</sub> *tkl1*Δ<br>CM<sub>TKL2</sub> *tkl1*Δ |
| *TRP1* | *trp1*Δ::nat<sup>R</sup> | *TRP1*<br>Φ<br>*trp1*<sup>C17A</sup><br>*trp1*<sup>C17M</sup> | WT<sub>TRP1</sub><br>KO<sub>TRP1</sub><br>CM.1<sub>TRP1</sub><br>CM.2<sub>TRP1</sub> |

<sup>a</sup> All strains were generated on the Y8205 parental strain (MAT-α *can1*Δ::STE2pr-Sp\_his5 *lyp1*Δ::STE3pr-LEU2 *his3*Δ *leu2*Δ *ura3*Δ)
<sup>b</sup> Centromeric low-copy number plasmids were from the MoBY-ORF collection, the empty vector is p5586 is p-Φ. Plasmid inserts include the yeast ORF with its native 5'- and 3'-UTR regions.

**Table S2.**
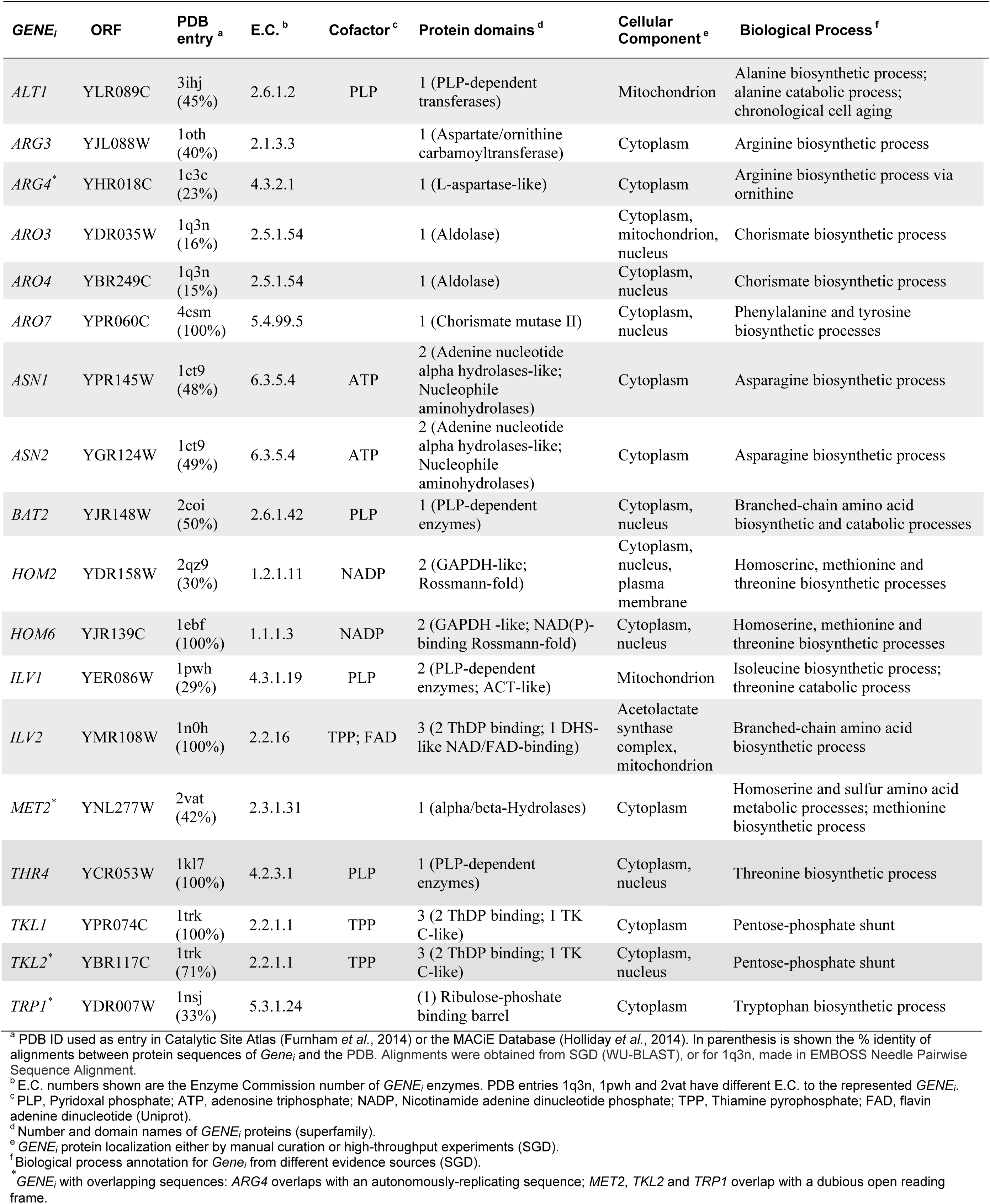
Complete information of amino acid biosynthesis enzymes in this study.

| *GENE<sub>i</sub>* | ORF | PDB entry <sup>a</sup> | E.C. <sup>b</sup> | Cofactor <sup>c</sup> | Protein domains <sup>d</sup> | Cellular Component <sup>e</sup> | Biological Process <sup>f</sup> |
| --- | --- | --- | --- | --- | --- | --- | --- |
| *ALT1* | YLR089C | 3ihj (45%) | 2.6.1.2 | PLP | 1 (PLP-dependent transferases) | Mitochondrion | Alanine biosynthetic process; alanine catabolic process; chronological cell aging |
| *ARG3* | YJL088W | 1oth (40%) | 2.1.3.3 |  | 1 (Aspartate/ornithine carbamoyltransferase) | Cytoplasm | Arginine biosynthetic process |
| *ARG4\** | YHR018C | 1c3c (23%) | 4.3.2.1 |  | 1 (L-aspartase-like) | Cytoplasm | Arginine biosynthetic process via ornithine |
| *ARO3* | YDR035W | 1q3n (16%) | 2.5.1.54 |  | 1 (Aldolase) | Cytoplasm, mitochondrion, nucleus | Chorismate biosynthetic process |
| *ARO4* | YBR249C | 1q3n (15%) | 2.5.1.54 |  | 1 (Aldolase) | Cytoplasm, nucleus | Chorismate biosynthetic process |
| *ARO7* | YPR060C | 4csm (100%) | 5.4.99.5 |  | 1 (Chorismate mutase II) | Cytoplasm, nucleus | Phenylalanine and tyrosine biosynthetic processes |
| *ASN1* | YPR145W | 1ct9 (48%) | 6.3.5.4 | ATP | 2 (Adenine nucleotide alpha hydrolases-like; Nucleophile aminohydrolases) | Cytoplasm | Asparagine biosynthetic process |
| *ASN2* | YGR124W | 1ct9 (49%) | 6.3.5.4 | ATP | 2 (Adenine nucleotide alpha hydrolases-like; Nucleophile aminohydrolases) | Cytoplasm | Asparagine biosynthetic process |
| *BAT2* | YJR148W | 2coi (50%) | 2.6.1.42 | PLP | 1 (PLP-dependent enzymes) | Cytoplasm, nucleus | Branched-chain amino acid biosynthetic and catabolic processes |
| *HOM2* | YDR158W | 2qz9 (30%) | 1.2.1.11 | NADP | 2 (GAPDH-like; Rossmann-fold) | Cytoplasm, nucleus, plasma membrane | Homoserine, methionine and threonine biosynthetic processes |
| *HOM6* | YJR139C | 1ebf<br>(100%) | 1.1.1.3 | NADP | 2 (GAPDH -like; NAD(P)-binding Rossmann-fold) | Cytoplasm, nucleus | Homoserine, methionine and threonine biosynthetic processes |
| *ILV1* | YER086W | 1pwh<br>(29%) | 4.3.1.19 | PLP | 2 (PLP-dependent enzymes; ACT-like) | Mitochondrion | Isoleucine biosynthetic process; threonine catabolic process |
| *ILV2* | YMR108W | 1n0h<br>(100%) | 2.2.16 | TPP; FAD | 3 (2 ThDP binding; 1 DHS-like NAD/FAD-binding) | Acetolactate synthase complex, mitochondrion | Branched-chain amino acid biosynthetic process |
| *MET2* * | YNL277W | 2vat<br>(42%) | 2.3.1.31 |  | 1 (alpha/beta-Hydrolases) | Cytoplasm | Homoserine and sulfur amino acid metabolic processes; methionine biosynthetic process |
| *THR4* | YCR053W | 1kl7<br>(100%) | 4.2.3.1 | PLP | 1 (PLP-dependent enzymes) | Cytoplasm, nucleus | Threonine biosynthetic process |
| *TKL1* | YPR074C | 1trk<br>(100%) | 2.2.1.1 | TPP | 3 (2 ThDP binding; 1 TK C-like) | Cytoplasm | Pentose-phosphate shunt |
| *TKL2* * | YBR117C | 1trk<br>(71%) | 2.2.1.1 | TPP | 3 (2 ThDP binding; 1 TK C-like) | Cytoplasm, nucleus | Pentose-phosphate shunt |
| *TRP1* * | YDR007W | 1nsj<br>(33%) | 5.3.1.24 |  | (1) Ribulose-phosphate binding barrel | Cytoplasm | Tryptophan biosynthetic process |

<sup>a</sup> PDB ID used as entry in Catalytic Site Atlas (Furnham *et al.*, 2014) or the MACiE Database (Holliday *et al.*, 2014). In parenthesis is shown the % identity of alignments between protein sequences of *Gene<sub>i</sub>* and the PDB. Alignments were obtained from SGD (WU-BLAST), or for 1q3n, made in EMBOSS Needle Pairwise Sequence Alignment.
<sup>b</sup> E.C. numbers shown are the Enzyme Commission number of *GENE<sub>i</sub>* enzymes. PDB entries 1q3n, 1pwh and 2vat have different E.C. to the represented *GENE<sub>i</sub>*.
<sup>c</sup> PLP, Pyridoxal phosphate; ATP, adenosine triphosphate; NADP, Nicotinamide adenine dinucleotide phosphate; TPP, Thiamine pyrophosphate; FAD, flavin adenine dinucleotide (Uniprot).
<sup>d</sup> Number and domain names of *GENE<sub>i</sub>* proteins (superfamily).
<sup>e</sup> *GENE<sub>i</sub>* protein localization either by manual curation or high-throughput experiments (SGD).
<sup>f</sup> Biological process annotation for *Gene<sub>i</sub>* from different evidence sources (SGD).
\* *GENE<sub>i</sub>* with overlapping sequences: *ARG4* overlaps with an autonomously-replicating sequence; *MET2*, *TKL2* and *TRP1* overlap with a dubious open reading frame.

**Table S3.**
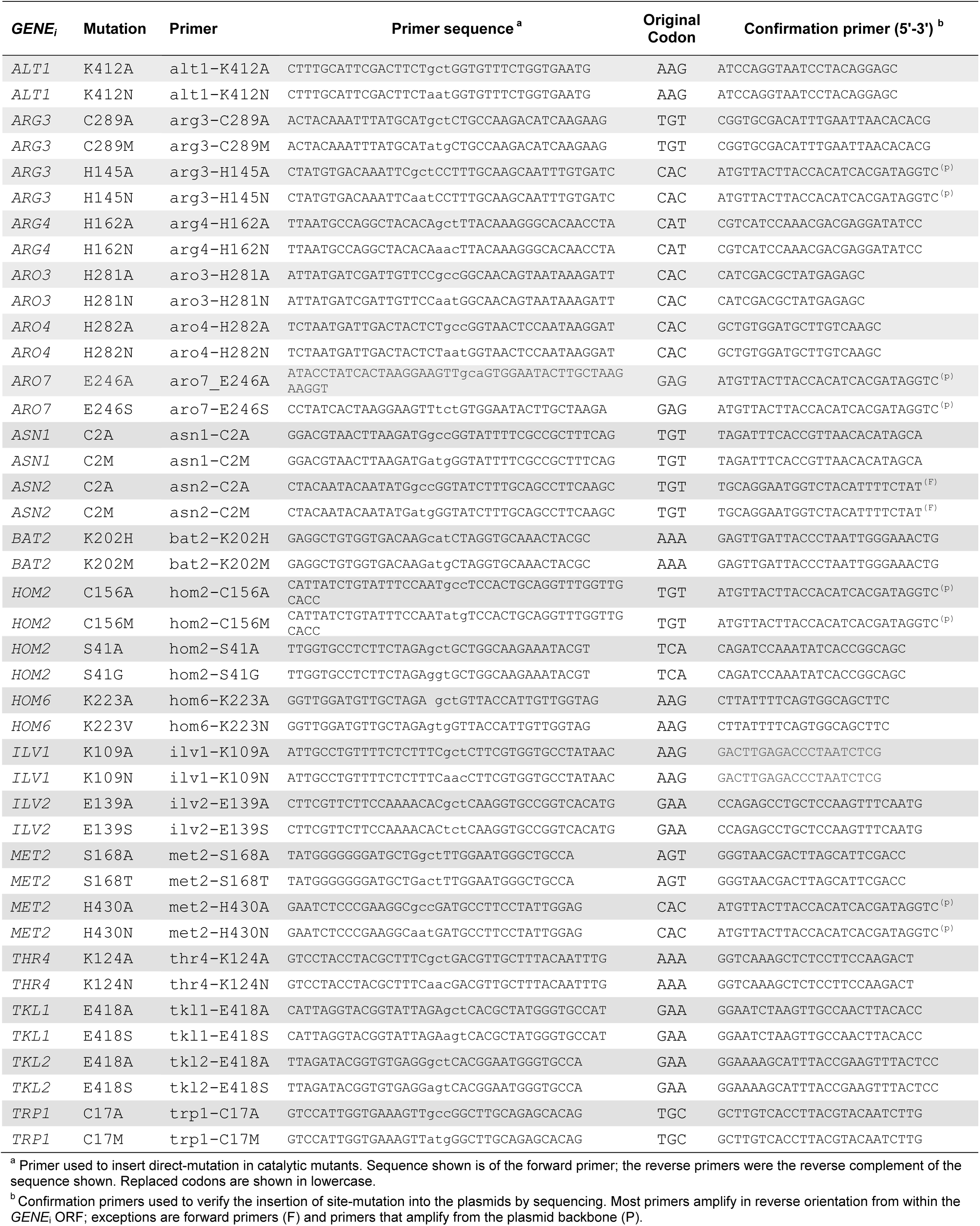
Primers used for site-directed mutagenesis.

**Table S4.**
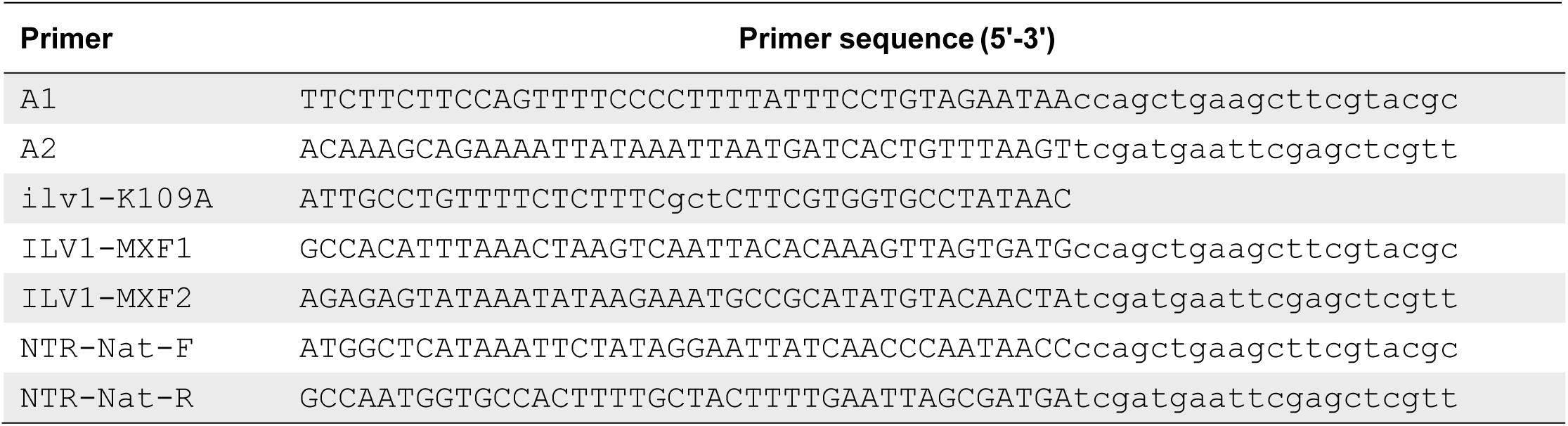
Primers used for genome-integrated mutations.

**Table S5.** Gene Ontology enrichment analysis of *ILV1* genetic interactions scored in this study.

| GO Term | Description <sup>a</sup> | P-value | FDR q-value | B<sup>b</sup> | b<sup>c</sup> |
| --- | --- | --- | --- | --- | --- |
| GO:0032543 | mitochondrial translation | 3.33E-8 | 1.45E-4 | 67 | 30 |
| GO:0043170 | macromolecule metabolic process | 8.57E-6 | 9.34E-3 | 1333 | 265 |
| GO:0044260 | cellular macromolecule metabolic process | 4.33E-6 | 9.44E-3 | 1253 | 253 |
| GO:0006325 | chromatin organization (6) | 6.68E-6 | 9.71E-3 | 177 | 52 |
| GO:0044237 | cellular metabolic process | 2.72E-5 | 1.7E-2 | 1860 | 349 |
| GO:0034645 | cellular macromolecule biosynthetic process (5) | 3.82E-5 | 1.85E-2 | 554 | 124 |
| GO:0006807 | nitrogen compound metabolic process | 2.45E-5 | 2.14E-2 | 1582 | 304 |
| GO:0031326 | regulation of cellular biosynthetic process (12) | 5.45E-5 | 2.38E-2 | 515 | 116 |
| GO:0006354 | DNA-templated transcription, elongation | 1.87E-4 | 5.1E-2 | 28 | 13 |
| GO:0044267 | cellular protein metabolic process | 1.78E-4 | 5.19E-2 | 610 | 131 |
| GO:0071704 | organic substance metabolic process | 2.39E-4 | 5.22E-2 | 1840 | 340 |
| GO:0044238 | primary metabolic process | 2.18E-4 | 5.27E-2 | 1752 | 326 |
| GO:0009058 | biosynthetic process | 2.76E-4 | 5.48E-2 | 879 | 178 |
| GO:0045721 | negative regulation of gluconeogenesis | 3.97E-4 | 6.66E-2 | 8 | 6 |
| GO:0016070 | RNA metabolic process | 5.37E-4 | 7.1E-2 | 531 | 114 |
| GO:0034641 | cellular nitrogen compound metabolic process | 5.35E-4 | 7.29E-2 | 1039 | 204 |
| GO:0070271 | protein complex biogenesis | 6.38E-4 | 7.73E-2 | 31 | 13 |
| GO:0019538 | protein metabolic process | 4.98E-4 | 7.76E-2 | 735 | 151 |
| GO:0043457 | regulation of cellular respiration | 7.2E-4 | 8.26E-2 | 4 | 4 |
| GO:0008152 | metabolic process | 7.09E-4 | 8.36E-2 | 2028 | 367 |
| GO:0000959 | mitochondrial RNA metabolic process | 8.25E-4 | 8.57E-2 | 21 | 10 |

<sup>1</sup> Non-redundant enriched biological process GO-terms (P-value < 10<sup>-3</sup>, FDR q-value < 0.1) from genetic interactions (|S-score| > 3) of both profiles from knockout (*ilv1Δ*) and catalytic mutant (*ilv1K109A*). Total genes= 3,399; Genetic interaction genes= 558.
<sup>a</sup> In parenthesis are the numbers of enriched GO terms not shown given a 50% redundancy to the corresponding entry.
<sup>b</sup> B is the total number of genes associated with a specific GO term.
<sup>c</sup> b is the number of gene-hits associated with a specific GO term.

**Table S6.** Gene Ontology enrichment analysis of the non-catalytic genetic interactions of *ILV1*.

| GO Term | Description | P-value | FDR q-value | B<sup>a</sup> | b<sup>b</sup> |
| --- | --- | --- | --- | --- | --- |
| GO:0005739 | mitochondrion | 2.39E-4 | 1.85E-1 | 459 | 54 |
| GO:0044429 | mitochondrial part | 3.37E-4 | 1.31E-1 | 260 | 35 |
| GO:0070271 | protein complex biogenesis | 7.58E-6 | 2.97E-2 | 12 | 7 |
| GO:0070823 | HDA1 complex | 4.36E-4 | 1.13E-1 | 3 | 3 |

<sup>1</sup> Non-redundant GO-terms from biological process and cellular component GO-terms (P-value < 10<sup>-3</sup>, FDR q-value < 0.2) from non-catalytic genetic interactions of ILV1 (S-score < -3; p<0.005). Total genes= 2,230; non-catalytic genetic interactors= 170.
<sup>a</sup> B is the total number of genes associated with a specific GO term.
<sup>b</sup> b is the number of gene-hits associated with a specific GO term.

**Dataset S1.** Phenotypic screen, complete data set (XLS).

**Dataset S2.** Differential epistasis profile analysis, complete data set (XLS).

